# Differentiation status determines tumorigenicity and immunogenicity of cancer cells

**DOI:** 10.1101/2025.05.30.656250

**Authors:** Yang Liu, Chen Wang, Jing Li, Ying Cao

## Abstract

Cancer (tumorigenic) cells exhibit cellular properties such as tumorigenicity, evasion of anti-tumor immunity and cell death, metastasis, etc. We showed previously that the core property of cancer cells is neural stemness, which represents general stemness and confers cancer cells with tumorigenicity and pluripotency. Immunogenicity of cancer cells plays the key role in anti-tumor immunity. While tumorigenicity is defined by neural stemness and its regulatory networks, how immunogenicity of cancer cells is determined and how tumorigenicity is related with immunogenicity remained unknown. In the present study, we show that genes conferring tumorigenicity and genes conferring immunogenicity are inversely correlated with differentiation status of cells. Cancer cells can be induced to differentiate into different types of cells either by lineage specific differentiation factors, e.g., muscle differentiation factor MYOD1 or adipocyte differentiation factor PPARG, or by blocking cell-intrinsic oncofactors, e.g., SETDB1. Induced differentiation reprograms both cellular properties and transcriptomes of cancer cells, leading to suppression of genes conferring tumorigenicity and upregulation of genes conferring immunogenicity, and ultimately, leading to loss of neural stemness, suppression of tumorigenicity and enhancement of immunogenicity in cancer cells after differentiation. Vice versa, dedifferentiation leads to the opposite effect. The results identified the **cellular mechanism** that tumorigenicity and immunogenicity of cancer cells are inversely connected and determined by differentiation status. Such a mechanism, together with our previous studies, reinforces that cancer cell properties should be understood by understanding neural stemness and the principle of embryonic cell/tissue differentiation, and novel cancer therapies can be developed by employing differentiation effect of cancer cells induced by differentiation factors. As a proof of concept test, we showed that muscle differentiation factor MYOD1 suppresses tumorigenesis in a mouse model of hepatocellular carcinoma.

## Introduction

Our previous studies demonstrated that cancer or tumorigenic cells are characteristic of the properties of and share regulatory networks with neural stem cells (NSCs), i.e., neural stemness, which represents the general stemness rather than a type of tissue stemness, and confers both tumorigenicity and pluripotency in cancer cells (Zhang et al., 2017; Lei et al., 2019; Chen et al., 2021; Xu et al., 2021; Cao, 2022; Zhang et al., 2022; Cao, 2023). Pluripotent property of cancer cells was first characterized in teratocarcinoma cells, which subsequently enlightened the research of pluripotency of embryonic stem cells (Solter, 2006). This means that cancer cells, like embryonic pluripotent cells, can be induced to differentiate into different types of cells (Cao, 2022; Cao, 2023). Cells of a type exhibit a variety of cellular properties specific to the cell type that are defined by cell-specific gene regulatory network, such as morphology, adhesiveness, mobility, proliferation, differentiation potential, metabolism, physiological function, etc. Therefore, distinct cell types should differ in their cellular properties and regulatory networks. For example, pluripotent embryonic cells and maturely differentiated cells, which are defined by different gene regulatory networks, show different cellular properties. Cancer cells are characteristic of high proliferation, migration, evasion of cell death and immune destruction, etc., which are defined by the gene regulatory network of embryonic neural or neural stem cells (Zhang et al., 2017; Cao, 2022). It can be deduced that the regulatory networks and properties of cancer cells will be reprogrammed if they can be induced to differentiate into a more mature form.

Cancer cells are capable of immunoevasion via complex mechanisms (Kim and Cho, 2022; Vinay et al., 2015). Immunotherapies have revolutionized cancer treatment, but achieve responses in only a minority of patients due to primary, adaptive or acquired resistance (Draghi et al., 2019; Gide et al., 2018; Kalbasi and Ribas, 2020; O’Donnell et al., 2019; Schoenfeld and Hellmann, 2020; Shah and Fry, 2019; Sharma et al., 2017; Vitale et al., 2021), and sometimes cause even hyperprogression of cancer (Champiat et al., 2018; de Miguel and Calvo, 2020; Kamada et al., 2019; Marcucci and Rumio, 2021). Inhibitors of immune checkpoints PD-1/PD-L1 or CTLA-4 play the major role in cancer immunotherapy. Clinical trials of immune checkpoints identified later, e.g., IDO1 and TIGIT, did not achieve favorable results, suggesting that it might be not sufficient to understand the functions of immune checkpoints in cancer merely in the context of immunity. The mechanisms for resistance to immunotherapy are also a complicated issue, but several possibilities have been proposed, including insufficient tumor antigenicity due to the lack of tumor neoantigens, defects in transduction of anti-tumor immune response mediated by tumour-intrinsic IFN-γ signaling, impaired antigen processing and presentation machinery, regulation by oncogenic signaling, and tumor dedifferentiation and stemness (Draghi et al., 2019; Kalbasi and Ribas, 2020; Schoenfeld and Hellmann, 2020; Sharma et al., 2017). In other words, these defects can be interpreted as the consequences of repression of genes involved in anti-tumor immune response or immunogenicity of cancer cells, for instances, by MEX3B (Huang et al., 2018), CDK4/6 (Deng et al, 2018), ADAR1 (Ishizuka et al., 2019), EZH2 (Kim et al., 2020; Zhou et al., 2020), β-catenin (Ruiz de Galarreta et al., 2019; Spranger et al., 2015), SETDB1 (Griffin et al., 2021), C-MYC (Zimmerli et al., 2022), KRAS (Watterson and Coelho, 2023; Lasse-Opsahl et al., 2025). Although the detailed molecular mechanisms responsible for repression of immune related genes by these factors are different, they are intrinsically interconnected together by that all these factors are enriched in embryonic neural cells. This means that they are components of the regulatory network of embryonic neural cells or NSCs. They are also well known oncoproteins that play extensive roles in cancer initiation and progression. However, “oncogenic signaling is the least understood aspect of functional immunogenicity” (Karasarides et al., 2022). Immunoevasion and immunotherapy resistance effect mediated by these embryonic neural/neural stemness factors should be a result of acquirement or enhancement of neural stemness, which endows cancer cells with tumorigenicity and pluripotency (Zhang et al., 2017; Xu et al., 2021; Cao, 2022; Zhang et al., 2022). In agreement, dedifferentiation and stemness of cancer cells is the key factor driving resistance to immunotherapy (Miao et al., 2019; Lei and Lee, 2021; Li and Stanger, 2020). Cancer-initiating cells exhibit immune privilege, protecting them from immune attack by making use of the mechanisms above and the expression of immune checkpoints (Galassi et al., 2021; Joyce and Fearon, 2015). NSCs also exhibit immune privilege, because they express low levels of immune-related proteins, including MHC class I and II antigens, HLA-DR and co-stimulatory molecules (Itakura et al., 2017; Magliocca et al., 2006; Ozaki et al., 2017). Moreover, blocking oncoproteins and immunotherapy resistance factors, such as EZH2, LSD1 or SETDB1, in cancer cells causes a neuronal differentiation effect and loss of tumorigenicity (Zhang et al 2017; Lei et al., 2019; Zhang et al., 2022). These results suggest that immunogenicity might be related with differentiation status of cancer cells. This relationship could also be deduced from developmental origin of immune related genes and neural stemness. Immune system is derived from mesodermal differentiation during embryonic development. Immune-related genes are primarily expressed in immune cells and non-neural tissues/organs. Therefore, expression of cancer promoting genes, most of which are embryonic neural/neural stemness genes (Zhang et al., 2017), and expression of immune related genes should be generally mutually exclusive, implying that tumorigenicity and immunogenicity are mutually exclusive properties. In the present study, we show a general tendency that expression of immune related genes is generally elevated in differentiated cells. Cancer cells can be induced to differentiate into different types of cells by lineage specific differentiation factors or by blocking cell-intrinsic oncofactors. Differentiation status of cancer cells is negatively correlated with tumorigenicity and positively correlated with immunogenicity because of general inverse correlation of expression between cancer promoting genes and immune related genes. Differentiation effects reprogram both cellular properties and transcriptomes of cancer cells, leading to loss of neural stemness, suppression of tumorigenicity and enhancement of immunogenicity in cancer cells after differentiation. These results help to clarify that the basic properties of cancer cells are intrinsically connected by neural stemness. We present the evidence that a lineage differentiation factor, MYOD1, inhibits tumorigenesis in a mouse model of hepatocellular carcinoma. We identified the **cellular mechanism** underlying the properties of cancer cells, particularly tumorigenicity and immunogenicity, and suggest that novel cancer therapies might be developed by employing differentiation effect of cancer cells induced by differentiation factors.

## Results

### Cancer cell differentiation by forced expression of MYOD1, PPARG, or knockdown of SETDB1

Cancer (tumorigenic) cells are characteristic of neural stemness and pluripotent different potential, we reasoned that key factors driving tissue differentiation during embryogenesis, such as muscle differentiation factor MYOD1 and adipocyte differentiation factor PPARG, could also drive cancer cell differentiation. Forced expression of MYOD1 via lentiviral transfection led to a significant morphological change in the HCT116 colorectal cancer cells, as compared with control cells that were transfected with virus carrying the empty vector (Ctrl) (Fig. 1A), and forced expression of PPARG in HCT116 cells caused adipocyte-like differentiation, as confirmed by oil-red staining (Fig. 1A). MYOD1 expression caused stimulation of muscle-specific protein MEF2C, and PPARG stimulated expression of adipocyte marker FABP4 (Fig. 1B), suggestive of muscle- or adipocyte-like differentiation in HCT116 cells, respectively. However, oncofactors SETDB1, EZH2, DNMT1, C-MYC, the proliferation marker PCNA, and neural stemness markers PAX6, SOX1, SOX2, and MSI1, all showing specific or enriched expression in embryonic neural cells, were repressed (Fig. 1B), suggesting that neural stemness, and malignant features and tumorigenicity in cells might be reduced in response to MYOD1 or PPARG expression. Indeed, control cells displayed neural stemness, which was manifested by formation of neurosphere-like structure in NSC-specific serum-free culture, as reported previously (Xu et al., 2021; Zhang et al., 2022). By contrast, cells with forced MYOD1 or PPARG expression failed to form neurospheres (Fig. S1A), indicating loss of neural stemness. Moreover, these cells showed decreased capability of colony formation in soft agar and reduced ability of migration and invasion (Fig. S1B-D). Xenograft tumor formation in immunodeficient nude mice is an indication of tumorigenicity. Tumors formed by cells with MYOD1 or PPARG expression were smaller than those formed by control cells (Fig. 1C), indicating suppressed tumorigenicity of differentiated cells. Accordingly, tumors formed by control cells showed stronger signals of C-MYC, SETDB1, and PCNA than those formed by cells with forced MYOD1 or PPARG expression. Moreover, the immune checkpoint PD-L1 was also more strongly detected in tumors of control cells (Fig. 1D). Oncoproteins like C-MYC and SETDB1 and immune checkpoints promote immune evasion. Therefore, their stronger expression in tumor suggests stronger capability of immune evasion. Cancer cells are capable of pluripotent differentiation, for example, neuronal-like differentiation (Zhang et al., 2017; Lei et al., 2019; Xu et al., 2021; Zhang et al., 2022). Neuronal marker MAP2 was highly detected in control tumors, indicating significant neuronal differentiation, but its signal was weak in tumors formed by cells with forced MYOD1 or PPARG expression (Fig. 1D). This implies that differentiation potential of HCT116 cells was curbed when they already differentiated, in agreement with the results reported previously (Zhang et al., 2022).

**Fig. 1.**
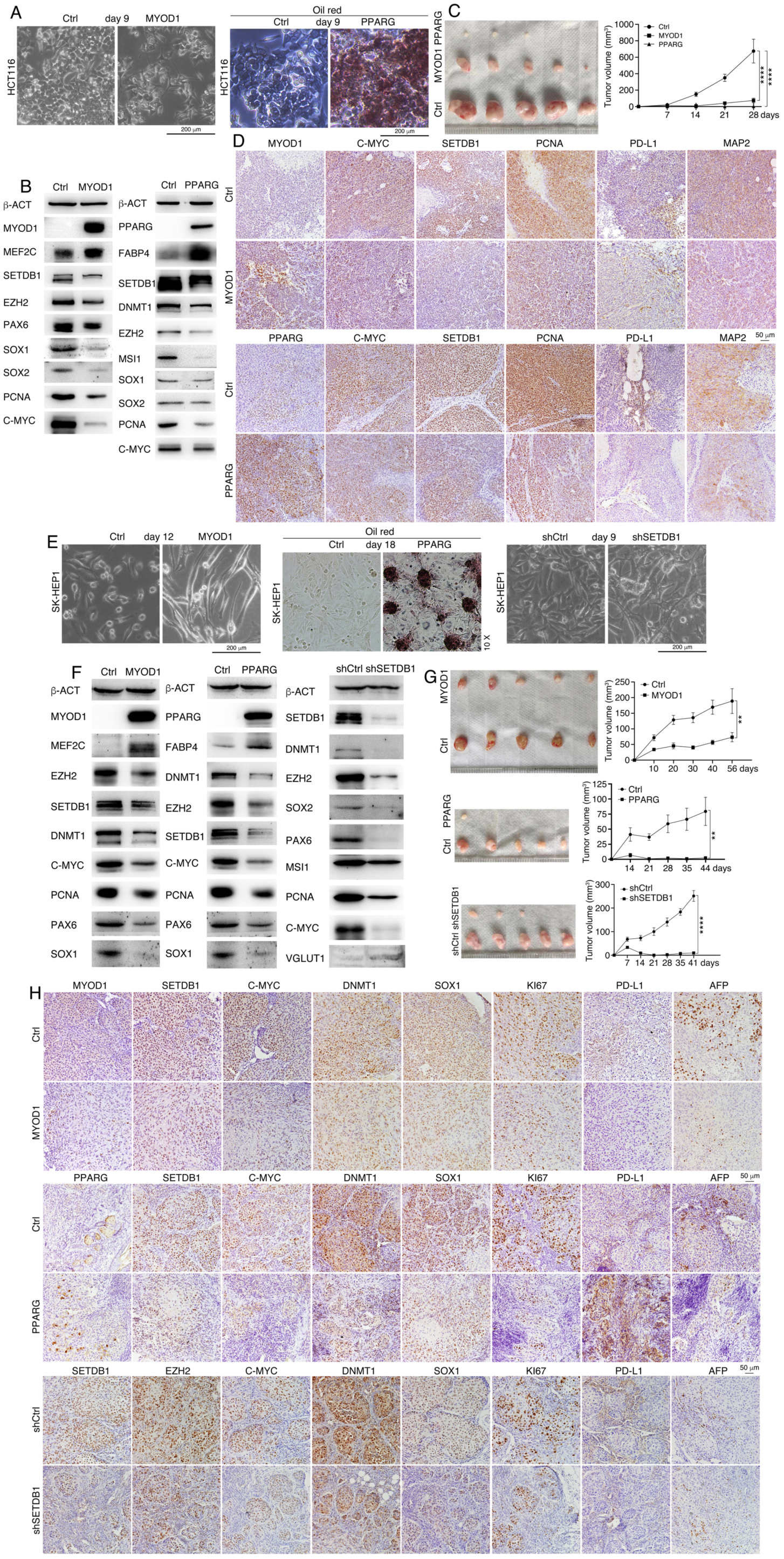
Characterization of the effect of induced differentiation of cancer cells induced by MYOD1 or PPARG expression or by SETDB1 knockdown. (A-D) Induced differentiation of HCT116 cells induced by MYOD1 or PPARG expression. (A) Phenotypic change in cells in response to forced expression of MYOD1 (left) or PPARG (right). Oil red staining was used to detect the presence of lipids in cells with forced PPARG expression. Cells transfected with lentivirus carrying empty vector were used as control (Ctrl). (B) WB detection of proteins marking muscle or adipocyte differentiation or cell proliferation, cancer promoting factors and neural stemness factors in control cells (Ctrl) and cells with forced expression of MYOD1 or PPARG. β-ACT was used as loading control. (C) Comparison of xenograft tumor formation by cells without (Ctrl) and with forced expression of MYOD1 or PPARG in nude mice. Significance of difference in tumor growth measured with tumor volume was calculated using two-way ANOVA-Bonferroni/Dunn test. Data are shown as mean ± SD. ****p < 0.0001. (D) IHC detection of protein expression in sections of xenograft tumors derived from control and treated cells. Objective magnification: 20 ×. (E-H) Induced differentiation of SK-HEP1 cells induced by MYOD1 or PPARG expression or by knockdown of SETDB1. (E) Phenotypic change in cells in response to forced expression of MYOD1 (left) or PPARG (middle) or knockdown of SETDB1 (right). Oil red staining was used to detect the presence of lipids in cells. Cells transfected with lentivirus carrying empty expression vector (Ctrl) or knockdown vector (shCtrl) were used as controls. (F) WB detection of proteins marking tissue-specific differentiation and cell proliferation, cancer promoting factors and neural stemness factors in control cells and cells with induced differentiation. β-ACT was used as loading control. (G) Comparison of xenograft tumor formation by control cells and cells with forced expression of MYOD1 or PPARG or knockdown of SETDB1 in nude mice. Significance of difference in tumor growth measured with tumor volume was calculated using two-way ANOVA-Bonferroni/Dunn test. Data are shown as mean ± SD. **p < 0.01, ****p < 0.0001. (H) IHC detection of protein expression in sections of xenograft tumors derived from control and treated SK-HEP1 cells. Objective magnification: 20 ×.

Forced MYOD1 or PPARG expression reprogrammed the transcriptome of HCT116 cells. In agreement with their respective roles in tissue differentiation, MYOD1 activity led to upregulation of muscle specific genes including *ACTA2*, *TNNI1*, *TNNC1*, *TNNT1*, and PPARG stimulated genes marking adipocytes or involved in adipogenesis, including *FABP4*, *PLIN4*, *SP100*, and *CEBPD*. Moreover, both MYOD1 and PPARG could upregulate transcription of a series of immune related genes, including IFN-γ response or antigen processing and presentation genes, such as *B2M*, *HLA-A*, *HLA-DMA*, *IFI27*, *IFIT1*. Importantly, more genes were upregulated than downregulated (Table S1). Gene ontology (GO) analysis showed that downregulated genes in differentiated cells induced by either MYOD1 or PPARG were mainly associated with DNA replication, DNA metabolic process, chromosome organization, etc. (Fig. S1E, F), suggesting decreased rate of cell cycle or proliferation. Upregulated genes differentiated cells induced by MYOD1 were mainly associated with immune response, antigen processing and presentation, proteolysis, and those upregulated by PPARG were enriched for translation, peptide metabolic process, lipid metabolic process, etc. (Fig. S1E, F), reflecting the correlation between upregulated genes and functional properties of HCT116 cells after differentiation. Gene set enrichment analysis (GSEA) also revealed that genes in differentiated cells induced by MYOD1 were enriched for gene sets of immune response and regulation of immune responses, including ‘allograft rejection’, ‘B cell receptor signaling pathway’, ‘Fc epsilon RI signaling pathway’, ‘graft versus host disease’, etc., (Fig. S1G), and genes in differentiated cells induced by PPARG were enriched for immune related gene sets such as ‘graft versus host disease’, ‘intestinal immune network for IgA production’, ‘PD-L1 expression and PD-1 checkpoint pathway in cancer’, and ‘TNF signaling pathway’ (Fig. S1H). The results indicate that MYOD1 or PPARG can induce muscle- or adipocyte-like differentiation in HCT116 cells, reprogram their transcriptomes and cellular properties, including malignant features and tumorigenicity, and might also change the property to induce immune response.

We tested whether forced expression of MYOD1 or PPARG could generate similar reprogramming effects in the hepatocellular carcinoma cell SK-HEP1. Moreover, we also tested whether differentiation and cellular property reprogramming effect could be achieved in SK-HEP1 via inhibition of SETDB1, an oncoprotein that is expressed primarily in embryonic neural cells and plays a role in maintaining neural stemness (Zhang et al., 2022). MYOD1 induced muscle cell-like phenotype in the cells, and oil-red staining showed that PPARG induced adipocyte differentiation in SK-HEP1 cells (Fig. 1E). Knockdown of SETDB1 with specific short-hairpin RNA (shSETDB1) led to an elongated morphology (Fig. 1E). MYOD1 and PPARG stimulated expression of muscle cell marker MEF2C and adipocyte marker FABP4, respectively, in SK-HEP1 cells. They both repressed expression of oncoproteins SETDB1, EZH2, DNMT1, C-MYC, the proliferation marker PCNA, and neural stemness markers PAX6 and SOX1 (Fig. 1F). Knockdown of SETDB1 also led to repression of DNMT1, EZH2, C-MYC, PCNA, and neural stemness proteins SOX2, PAX6, and MSI1, but neuronal protein VGLUT1 was upregulated (Fig. 1F). These data supported that differentiation effects occurred with possibly reduced neural stemness and malignant properties in cells. SK-HEP1 cells formed cell clusters but not free floating neurospheres in NSC-specific serum-free culture, which suggest weaker neural stemness (Xu et al., 2021). Forced MYOD1 or PPARG expression or SETDB1 knockdown reduced their capability of cluster formation (Fig. S2A), as well as the ability of colony formation (Fig. S2B), invasion and migration (Fig. S2C). Forced expression of MYOD1 or PPARG or SETDB1 knockdown reduced tumorigenicity of cells, as shown by smaller tumor formation in nude mice (Fig. 1G). In tumors formed by control cells, more intense signals of SETDB1, C-MYC, DNMT1, SOX1, and KI67 were detected, as compared with tumors formed by cells with forced expression of MYOD1 or PPARG or knockdown of SETDB1 (Fig. 1H). PD-L1 was significantly detected in control tumors, but weakly in tumors formed by forced expression of MYOD1 or SETDB1 knockdown. But it was expressed more strongly in tumors formed by differentiated cells induced by PPARG (Fig. 1H). The general difference in expression of proteins suggest that cells in tumors formed by cells with by forced expression of MYOD1 or PPARG or SETDB1 knockdown might exhibit weaker ability of evasion of anti-tumor immunity. Besides, similar difference in the signal for the endodermal differentiation marker AFP was observed (Fig. 1H), suggesting the differentiation potential of SK-HEP1 was reduced after induced differentiation.

Transcriptome assays showed upregulation of muscle (e.g., *TNNI1*, *TNNC1*, *MSTN*, *MYL1*), adipocyte specific genes (e.g., *FABP4*, *PLIN2*, *PLIN4*) or neuronal genes (e.g., *MAP2*, *SLC17A7*, *CELF5*) in response to MYOD1 or PPARG expression or SETDB1 knockdown, respectively. Meanwhile, a series of IFN-γ response genes, including *BST2*, *IFIH1*, *IFIT1*, *IRF1*, *STAT1*, were activated in response to MYOD1 or PPARG activity, or genes such as *HLA-B*, *HLA-DQA1*, *ICAM1*, *IFI27*, were activated in SETDB1 knockdown cells (Table S1). As observed in HCT116 cells, there was a general tendency that more immune related genes were upregulated than those downregulated in SK-HEP1 cells after differentiation (Table S1). GO analysis demonstrated that downregulated genes due to MYOD1-induced differentiation were primarily associated with sex differentiation, reproductive structure development, and blood vessel morphogenesis, and upregulated genes were associated with defense response to virus/other organisms, myotube differentiation, and striated muscle cell differentiation (Fig. S2D). In differentiated cells induced by PPARG, downregulated genes were linked with antigen processing and presentation, immune system process and immune response, and upregulated genes were linked with lipid transport, lipid metabolic process and immune response (Fig. S2E). Both up- and downregulated genes being associated with immune response might be due to the fact that SK-HEP1 cells are less tumorigenic, and some immune related genes are still expressed at relatively high level in the cells. But still, more IFN-γ response genes were activated by PPARG than those dowregulated (Table S1). In cells with SETDB1 knockdown, downregulated genes were associated with DNA replication, DNA metabolic process and chromosome organization that are involved in cell cycle or proliferation, and those upregulated were linked with immune system process, immune response, proteolysis, etc. (Fig. S2F). In addition, GSEA revealed that genes in differentiated cells induced by MYOD1 or PPARG or by loss of SETDB1 function were enriched in immune related gene sets including ‘Fc epsilon RI signaling pathway’, ‘PD-L1 expression and PD-1 checkpoint pathway in cancer’, ‘Toll-like receptor signaling pathway’, ‘T cell receptor signaling pathway’, ‘TNF signaling pathway’ (Fig. S2G-I). The results demonstrated that either forced expression of lineage differentiation factors MYOD1 or PPARR or blocking an endogenous oncoprotein SETDB1 can induce differentiation of cancer cells along different lineages, and reprogram their cellular properties and transcriptomes. Interestingly, differentiation effects cause a general tendency of upregulation of immune related genes, raising the question whether immune related gene expression is associated with differentiation status of cells, and whether differentiation status of cancer cells determines not only their tumorigenicity but also immunogenicity.

### Immune related gene expression is correlated with differentiation status of cells

Immune response relies on expression of immune related genes. Expression pattern data demonstrated that immune related genes are primarily expressed in non-neural cells. For instance, IFN-γ response genes, genes involved in antigen processing and presentation, and genes responsible for TNF-α signaling mediated cell death, *Casp8*, *Ctse*, *Gsdmd*, *Gzma*, *H2-Aa*, *H2-DMa*, *H2-T22*, *Icam1*, *Ifi27*, *Ifi30*, *Ifit1*, *Ifnar2*, *Ifngr1*, *Irf1*, *Psmb8, Ripk3,* and *Stat1*, are mainly transcribed in thymus and a broad range of other non-neural tissues/organs, but not or only weakly in neural tissues in E14.5 mouse embryos (Fig. S3). Interferon signaling, antigen presentation and TNF-α signaling are major pathways regulating cancer cell immunogenicity (Kearney et al., 2018). Likewise, *Myod1* is transcribed in musculature, diaphragm and in the mesenchyme of different tissues/organs, and *Pparg* transcription is detected in musculature, urethra and bladder fundus (Fig. S3). It can be seen that expression patterns of *Myod1* or *Pparg* overlap with those of immune related genes. This suggests a positive regulatory relationship between tissue differentiation genes and immune related genes, which is supported by the data obtained from cancer cell differentiation above. By contrast, *Setdb1* is enriched in neural tissues in mouse embryo (Fig. S3). The dedifferentiation genes *Sox2* and *Oct4* (*Pou5f1*) are well known for their specific expression in the inner cell mass, embryonic neural cells and primordial germ cells (Rathjen et al., 2002; Lee et al., 2010), but not in differentiated cells. These distinct expression patterns already imply a general negative regulatory relationship between genes in embryonic neural cells and genes in non-neural cells. Such as a relationship reflects the general rule that genes specifying the properties and functions of different types of cells should antagonize mutually to guarantee the identity and function of a type of cells and establish boundaries between different tissues and organs.

We analyzed transcriptomes of several types of cells representing undifferentiated and differentiated status, i.e., embryonic stem cells derived from C57BL/6J (B6) mouse (mESCs), primitive neural stem cells (primNSCs) derived from mESCs, fat cells, muscle cells, and cells from spleen and thymus of 4-week B6 mouse. GO analyses were made by using the transcriptome of primNSCs as reference because the default state of ESCs is NSCs, and neural stemness represents the general stemness (Cao, 2022). In mESCs, genes with decreased transcription were associated primarily with ribosome biogenesis, ncRNA and rRNA processing, but genes with increased transcription were linked with angiogenesis and regulation vasculature development (Fig. 2A, B). This is in agreement with the notion that ESCs are primed with the potential of non-neural differentiation by TGF-β signaling (Cao, 2022), which plays the key role in maintaining ESC identity. In both muscle and fat cells, genes with decreased transcription were associated with ncRNA metabolic process, ribonucleoprotein complex biogenesis, ribosome biogenesis, RNA splicing, etc. (Fig. 2C, E), whereas the increased genes are associated with angiogenesis, muscle tissue development, fatty acid metabolic process, leukocyte differentiation, and regulation of defense response (Fig. 2D, F). Transcription of genes involved in morphogenesis of an epithelium, ribosome biogenesis, ribonucleoprotein complex biogenesis, rRNA processing, etc., were similarly decreased in both spleen and thymus cells (Fig. 2G, I), and reasonably, transcription level of genes involved in immune response was increased (Fig. 2H, J). A general tendency can be found that genes involved in ribosome biogenesis and RNA processing/splicing are less transcribed in differentiated cells and even in ESCs as compared with primNSCs, and genes associated with cell-type specific properties are more transcribed. This is in well agreement with the notion that basic cellular machineries such as ribosome biogenesis work concertedly together to define a fast proliferative and pluripotent state in NSCs and cancer cells, and differentiation causes downregulation of these basic machineries including ribosome biogenesis (Chen et al., 2021; Cao, 2022). Accordingly, differentiated cells show low or no proliferation and differentiation potential.

**Fig. 2.**
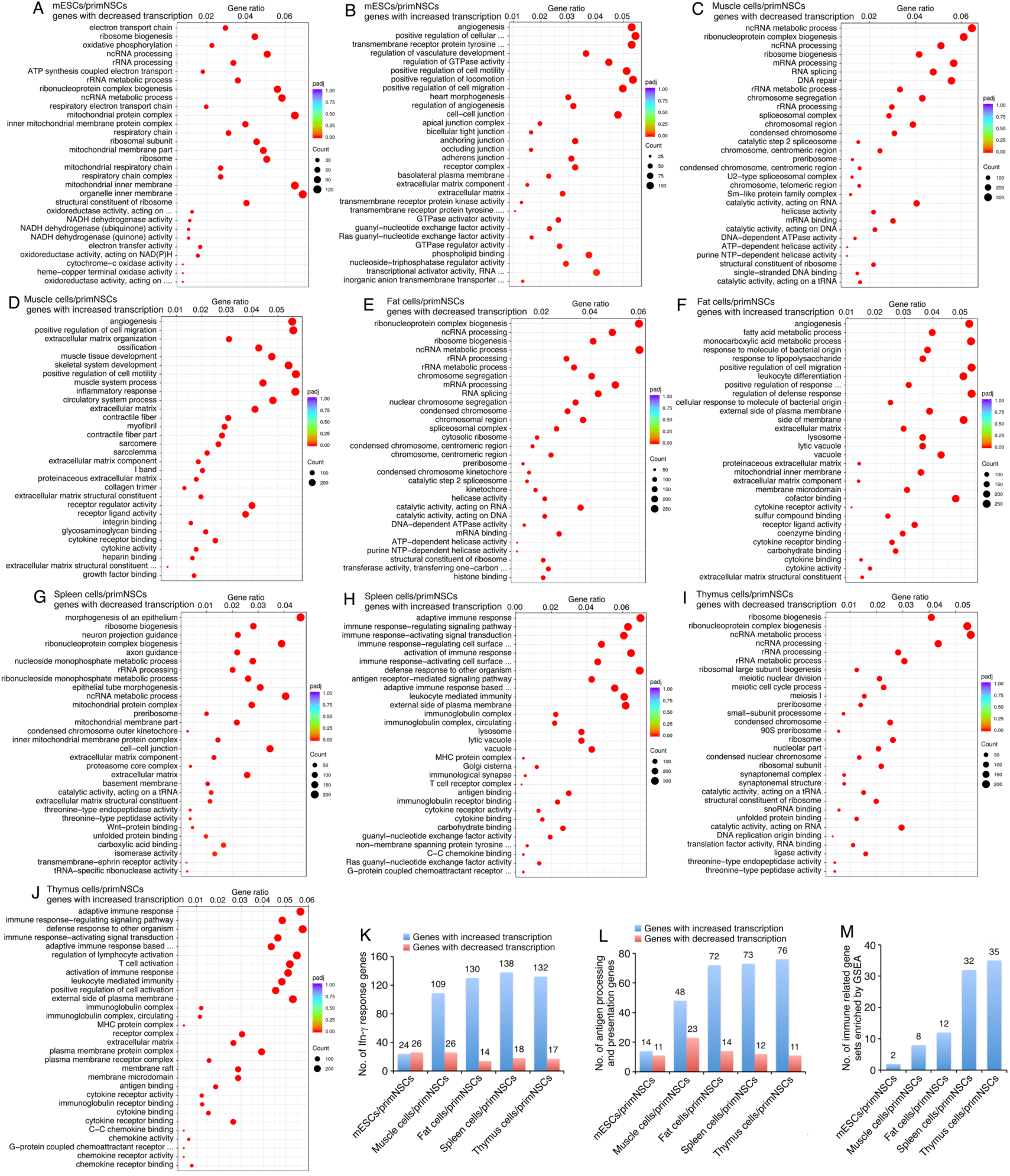
Transcriptome analysis on undifferentiated and differentiated cell types and correlation of immune related gene expression with differentiation status of cells. (A-J) GO enrichment of genes with decreased transcription (A, C, E, G, I) or genes with increased transcription (B, D, F, H, J) in the transcriptomes of mESCs, muscle cells, fat cells, spleen cells and thymus cells as compared to the transcriptome of primNSCs. (K-M) Difference in transcription of immune related genes between undifferentiated and differentiated cell types. Number of Ifn-γ response genes (K), number of antigen processing and presentation genes (L) that are differentially transcribed between undifferentiated and differentiated cell types, and the number of immune related gene sets enriched by GSEA (M) were counted.

We next analyzed whether there is also certain trend of change in transcription of immune related genes in these cells. Among 188 mouse Ifn-γ response genes, 24 showed increased transcription in mESCs and 26 were decreased. By contrast, 109 showed an increase in transcription and only 26 showed decrease in muscle cells. Similarly, many more Ifn-γ response genes were increased in transcription than those decreased in fat, spleen and thymus cells (Fig. 2K; Table S2). Among 129 antigen processing and presentation genes, transcription of 14 were increased but 11 were decreased in mESCs, 48 were increased and 23 decreased in muscle cells, and many more increased than decreased in fat, spleen and thymus cells (Fig. 2L; Table S2). GSEA revealed that genes in mESCs were enriched in two immune related gene sets, but enriched in 8, 12, 32, and 35 immune gene sets in muscle, fat, spleen and thymus cells, respectively (Fig. 2M; Table S2). These data, together with the observation on cancer cells above, demonstrated that in addition to the elevated level of transcription of cell-type relevant genes, differentiated cells express higher level of immune related genes, which suggests that they might exhibit higher immunogenicity.

### Induced differentiation of cancer cells reduces their tumorigenicity but enhances immunogenicity

We explored next whether differentiation of cancer cells via forced expression of Myod1 or Pparg or Setdb1 knockdown might change their cellular properties including immunogenicity. Myod1 activity in the melanoma cell line B16F10, which was derived from B6 mouse, caused dramatic phenotypic change, Pparg enabled oil-red staining, and knockdown of Setdb1 led to neurite-like outgrowth in cells (Fig. 3A). Myod1 and Pparg stimulated expression of muscle-specific protein Myog and adipocyte marker Fabp4, respectively, and neuron markers Syn1 and Map2 were upregulated in response to Setdb1 knockdown (Fig. 3B). Interestingly, either forced expression of Myod1 or Pparg or Setdb1 knockdown repressed oncoproteins Dnmt1, Ezh2, Setdb1, c-Myc, proliferation marker Pcna, and repressed neural stemness markers Pax6 and Sox1 (Fig. 3B), which suggested that differentiation towards different lineages led to loss of neural stemness and malignancy. It was confirmed by failure of neurospheric structure formation in NSC-specific serum-free culture and by reduced ability in colony formation, invasion and migration of cells upon differentiation (Fig. S4A-C). It was noteworthy that these oncoproteins not only promote tumorigenesis but also promote immune evasion or inhibit anti-cancer immunity. Transcriptome profiling demonstrated that genes downregulated in response to Myod1 activity were primarily associated with chromosome segregation, DNA replication and cell division, etc., and those upregulated associated with muscle differentiation and function (Fig. S5A). Pparg activity inhibited transcription of genes that are mainly involved in melanin or pigment metabolism in B16F10 cells, whereas it enhanced transcription of genes involved in immune responses (Fig. S5B). In cells with blocking of Setdb1, genes involved in DNA replication and cell division was downregulated, but those involved in regulation of ion transport and response to bacterium were upregulated (Fig. S5C). Muscle-specific genes such as *Myl1*, *Tnni1*, *Myog*, adipocyte-specific genes such as *Fabp4*, *Plin1*/*4*, or neuronal genes such as *Syn1*, *Tubb4a*, and *Sv2a*, were upregulated by Myod1 or Pparg expression or by Setdb1 knockdown, respectively, adding further support for differentiation towards different lineages. Besides, 22, 46 and 34 Ifn-γ response genes were upregulated with 18, 2 and 15 were downregulated in cells with forced Myod1 or Pparg expression or with Setdb1 knockdown, respectively. In the case of antigen processing and presentation genes, 7, 11 and 18 were increased, and 8, 1, and 7 were decreased, separately (Table S3). GSEA showed that genes in cells with Myod1 or Pparg expression or with Setdb1 knockdown were enriched for 6, 7, and 10 immune gene sets, such as ‘mature B cell differentiation involved in immune response’, ‘positive regulation of lymphocyte differentiation’, ‘regulation of natural killer cell activation’ (Fig. S5D-F). The result also suggests the trend that immune related genes are activated upon differentiation, which might change the immunogenicity of B16F10 cells.

**Fig. 3.**
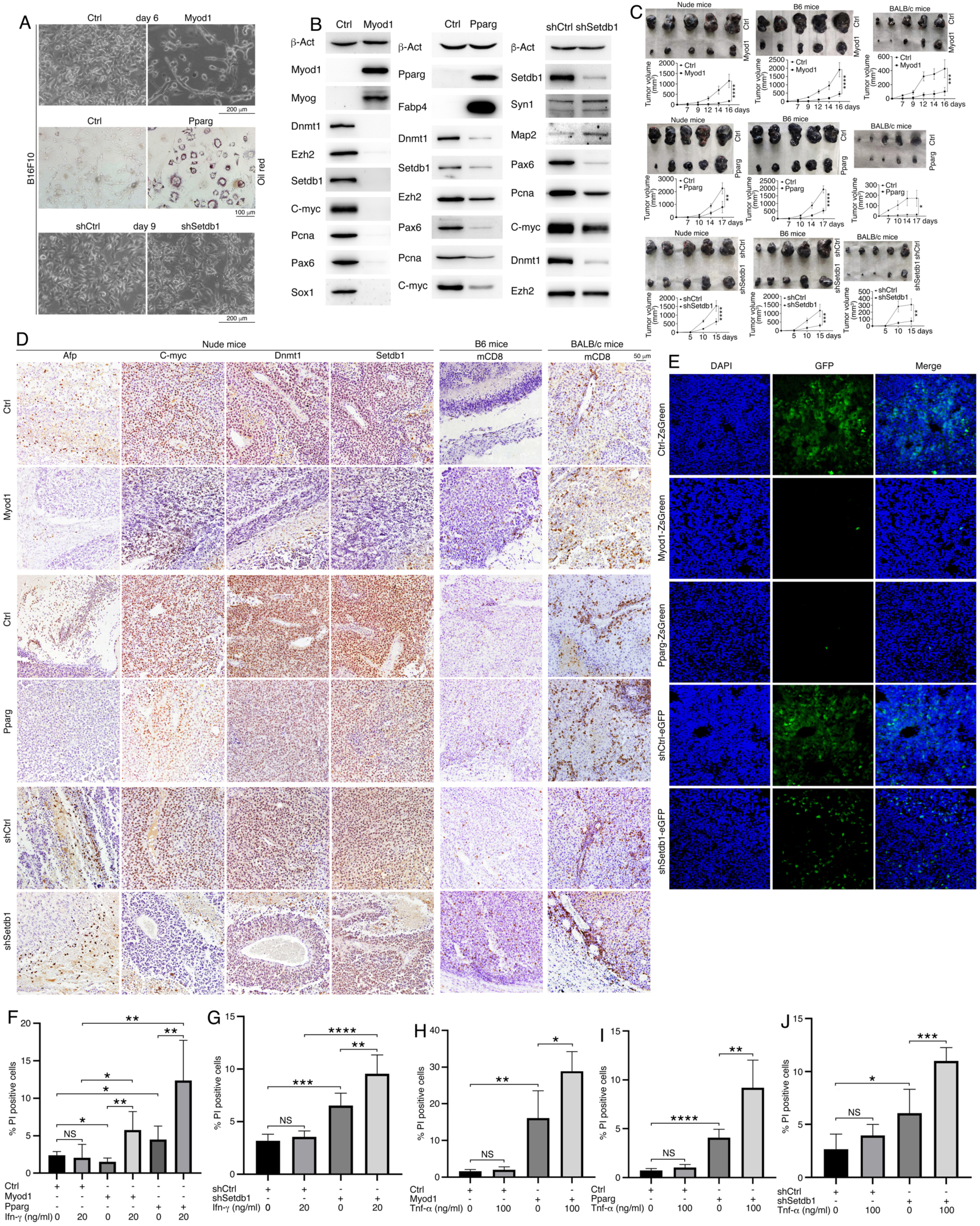
Induced differentiation reprograms cellular properties of B16F10 cells. (A) Induced differentiation effect in response to forced expression of Myod1 (upper) or Pparg (middle), or knockdown of Setdb1 (lower). Oil red staining (middle) was used to detect lipids. (B) WB detection of proteins marking lineage specific differentiation, cell proliferation, and neural stemness, and proteins that promote cancer. β-act was used as loading control. (C) Tumor formation by control cells and differentiated cells induced by Myod1, Pparg or inhibition of Setdb1 in immunodeficient (nude), syngeneic and allogeneic mice, respectively. Significance of difference in tumor growth measured with tumor volume was calculated using two-way ANOVA-Bonferroni/Dunn test. Data are shown as mean ± SD. *p < 0.05, **p < 0.01, ***p < 0.001, ****p < 0.0001. (D) IHC detection of protein or marker expression in sections of tumors formed by control and treated cells. Objective magnification: 20 ×. (E) Fluorescence in cryosections showing GFP-expressing cells in syngeneic tumors formed in B6 mice by B16F10 cells transfected with virus derived from ZsGreen-flagged vectors. (F-J) Difference in sensitivity of control cells and cells with induced differentiation to Ifn-γ (F, G) or Tnf-α (H-J) mediated cell death. Significance of difference was calculated using unpaired Student’s *t*-test. Data are shown as mean ± SD. *p < 0.05, **p < 0.01, ***p < 0.001, ****p < 0.0001. NS, not significant.

We then transplanted cells into nude, B6 and Balb/c mice, respectively, to analyze the effect of induced differentiation on tumorigenicity and immunogenicity of B16F10 cells. Cells with forced expression of Myod1, Pparg or knockdown of Setdb1 consistently formed smaller tumors than control cells in nude, B6 or Balb/c mice (Fig. 3C). Noteworthy is that syngeneic tumors were bigger than allogeneic tumors. We observed that xenograft tumors formed by control cells in nude mice exhibited stronger expression of an endodermal lineage differentiation marker Afp and oncoproteins C-myc, Dnmt1 and Setdb1 (Fig. 3D), an effect similar to those observed for HCT116 or SK-HEP1 cells. CD8+ signal, which represents T-cell infiltration in tumors, could be rarely detected in syngeneic tumors formed in B6 mice by control B16F10 cells, slightly stronger CD8+ signal was present in syngeneic tumors formed by cells with forced expression of Myod1 or Pparg or knockdown of Setdb1 (Fig. 3D). Substantially more CD8+ cells were detected in allogeneic tumors formed in Balb/c mice than in syngeneic tumors. There was also a general tendency that CD8+ signal was stronger in tumors formed by treated cells than in control tumors. It could be seen that difference in CD8+ signals was inversely correlated with the size of tumors formed by control and treated cells, and formed in syngeneic and allogeneic environment. Cryosections revealed that most cells in tumors formed by cells transfected with lentivirus containing the GFP-tagged empty vector were GFP-expressing cells. By contrast, very few or only a small portion of cells in tumors formed by cells transfected with lentivirus containing GFP-tagged Myod1 or Pparg expression vector or Setdb1 knockdown vector were GFP-expressing cells (Fig. 3E). This suggests that most cells with forced expression of Myod1 or Pparg or inhibition of Setdb1 did not contribute to tumor formation. Because of the general tendency of upregulation of genes involved in immune response in differentiated cancer cells, and Ifn-γ and Tnf-α signaling are the major pathways mediating immunogenicity of cancer cells (Kearney et al., 2018), we tested whether cancer cells altered their sensitivity to Ifn-γ and Tnf-α mediated cell death after differentiation. Control cells showed no significant change in sensitivity to Ifn-γ treatment, because no significant change in the percentage of propidium iodide (PI) positive cells, which marks dead cells. By contrast, when cells with forced expression of Myod1 or Pparg were treated with Ifn-γ, there was significant increase in the ratio of PI-positive cells (Fig. 3F; Fig. S6A). Increase in the ratio of PI-positive cells was also observed for Setdb1 knockdown cells after Ifn-γ treatment (Fig. 3G; Fig. S6A). Likewise, Tnf-α treatment of control cells did not cause significant change in the ratio of PI-positive cells, but treatment of cells with Myod1 or Pparg expression or Setdb1 knockdown led to increased ratio of PI-positive cells (Fig. 3H-J; Fig. S6B). These data indicate that induced differentiation enables B16F10 cells more sensitive to Ifn-γ and Tnf-α induced cell death, an effect accounting for the severely compromised contribution of differentiated cells to tumor formation.

We also tested the differentiation effect and the change in cellular properties in the CT26 colon cancer cells that were derived from Balb/c mouse. As observed in other cancer cells above, either forced expression of Myod1 or Pparg or inhibition of Setdb1 activity caused phenotypic changes in CT26 cells, and oil red staining could be seen in cells with Pparg expression (Fig. 4A). Besides stimulation of muscle protein Mef2c, adipocyte protein Fabp4, or neuronal proteins Map2 and Neun in these cells, proteins promoting cancer and immune evasion, and proteins marking neural stemness, were commonly repressed (Fig. 4B), suggestive of differentiation effects, loss of neural stemnes and malignancy. Indeed, the ability of colony formation, invasion and migration of cells after differentiation was inhibited (Fig. S4D-F). Myod1 caused repression of gene transcription associated mainly with cell adhesion, chemotaxis, blood vessel development, etc., and enhancement in transcription of genes associated with muscle contraction, nucleosome assembly, etc. (Fig. S7A). Pparg repressed transcription of genes involved in connective tissue, cartilage and muscle development, but upregulated genes mainly associated with innate immune response and responses to interferon β and γ (Fig. S7B). Setdb1 knockdown led to repression of genes associated with ribosome biogenesis and anion transport, and stimulation of genes associated with immune responses (Fig. S7C). Specifically, muscle, adipocyte or neuron genes were found among the upregulated genes in response to Myod1 or Pparg activity or Setdb1 knockdown, respectively, e.g. *Myl1*, *Tnni1*, *Myog* (muscle), *Fabp4*, *Plin2*/*4* (adipocyte), or *Eno2*, *Nefl*, *Sv2a* (neuron). In addition, 20, 31 and 55 Ifn-γ response genes were upregulated, and 20, 9 and 5 were downregulated, respectively (Table S3). A few antigen processing and presentation genes were changed in differentiated CT26 cells induced by Myod1 or Pparg, but 18 were upregulated with only 4 downregulated in cells with Setdb1 knockdown (Table S3). GSEA showed that genes in these cells were enriched for 5, 8, and 23 immune related gene sets, such as ‘antigen presentation via MHC class Ib’, ‘antigen processing and presentation’, or ‘myeloid cell activation involved in immune response’ (Fig. S7D-F).

**Fig. 4.**
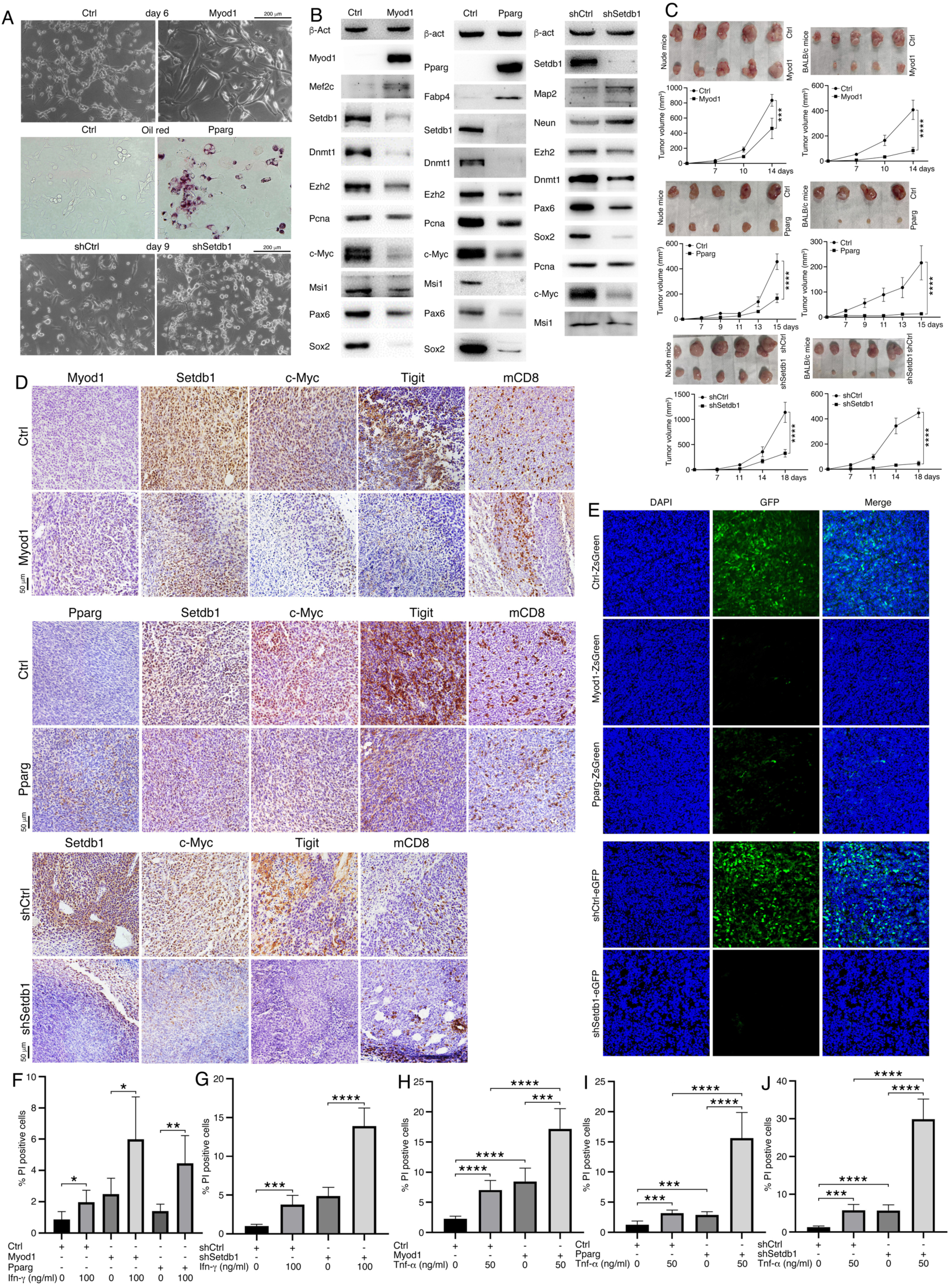
Induced differentiation reprograms cellular properties of CT26 cells. (A) Induced differentiation of CT26 cells by forced expression of Myod1 (upper) or Pparg (middle), or knockdown of Setdb1 (lower). Oil red staining (middle) showed presence of lipids. (B) WB detection of proteins marking lineage specific differentiation, cell proliferation, and neural stemness, and proteins that promote cancer. β-act was used as loading control. (C) Tumor formation by control cells and differentiated cells induced by Myod1, Pparg or inhibition of Setdb1 in immunodeficient (nude) and syngeneic mice, respectively. Significance of difference in tumor growth measured with tumor volume was calculated using two-way ANOVA-Bonferroni/Dunn test. Data are shown as mean ± SD. ***p < 0.001, ****p < 0.0001. (D) IHC detection of protein or marker expression in sections of tumors formed by control and treated cells. Objective magnification: 20 ×. (E) Fluorescence in cryosections showing GFP-expressing cells in syngeneic tumors formed in Balb/c mice by CT26 cells transfected with virus derived from ZsGreen-flagged vectors. (F-J) Difference in sensitivity of control cells and cells with induced differentiation to Ifn-γ (F, G) or Tnf-α (H-J) mediated cell death. Significance of difference was calculated using unpaired Student’s *t*-test. Data are shown as mean ± SD. *p < 0.05, **p < 0.01, ***p < 0.001, ****p < 0.0001. NS, not significant.

CT26 cells with forced expression of Myod1 or Pparg or Setdb1 knockdown led to reduced tumor formation in nude and syngeneic Balb/c mice (Fig. 4C), showing compromised tumorigenicity and enhanced immunogenicity of cells after differentiation. No tumor formation by either control or treated cells was observed in allogeneic B6 mice, suggesting that CT26 cells are more sensitive to immune rejection. It was found that oncoproteins Setdb1 and c-Myc, and the immune checkpoint Tigit, which inhibits both T and NK cell function in anti-cancer immunity, were more strongly detected in control tumors than in tumors formed by treated cells in syngeneic Balb/c mice, although no significant difference in CD8+ signal was observed (Fig. 4D). GFP labeling revealed that most cells with Myod1 or Pparg expression or Setdb1 knockdown didn’t contribute to tumor formation, either (Fig. 4D). Accordingly, differentiated CT26 cells induced by Myod1 or Pparg or by knockdown of Setdb1 were more sensitive to induced cell death when treated with Ifn-γ (Fig. 4F, G; Fig. S6C) or Tnf-α (Fig. 4H-J; Fig. S6D). In summary, induced differentiation of cancer cells reprograms their cellular properties and transcriptomes, causes a general tendency of upregulation of immune related genes, and consequently, reduces tumorigenicity but enhances immunogenicity of cancer cells.

### Dedifferentiation causes enhancement of tumorigenicity but reduction of immunogenicity in cancer cells

Mouse Oct4 and Sox2 and their homologous proteins in other species including human are well known for their capability of dedifferentiating differentiated cells into pluripotent state. Forced expression of Oct4 and SOX2 was performed to achieve a more dedifferentiated effect in CT26 cells via lentiviral transduction. Co-expression of Oct4 and SOX2 in CT26 cells (Oct4/SOX2 cells hereafter) led to earlier formation of neurospere-like structure than control cells in NSC-specific serum-free medium (Fig. 5A), an indication of enhancement of neural stemness. In addition, expression of neural stemness markers and cancer promoting factors was generally increased except c-Myc (Fig. 5B). Decreased expression of c-Myc might be due to that, during embryonic neural development, c-Myc is specifically expressed in neural crest cells and Oct4 and Sox2 are mainly expressed in neural plate, which gives rise to neural crest. Enhancement of neural stemness was accompanied with increased ability of cell invasion/migration and colony formation (Fig. S8A). RNAseq assay demonstrated that downregulated genes in dedifferentiated cells were mainly enriched in immune responses, and upregulated genes were enriched in interleukin-1-mediated signaling pathway and neural development (Fig. S8B), such as axon guidance. Interleukin-1 signaling and axon guidance are well characterized for their roles in promoting cancer progression (Jurcak et al., 2019; Garlanda and Mantovani, 2021; Lupo et al., 2024). Contrary to differentiation, dedifferentiation led to more Ifn-γ response genes downregulated than those upregulated, although no significance change in genes involved in antigen processing and presentation (Table S3). Oct4/SOX2 cells formed larger tumors in both nude mice and syngeneic Balb/c mice (Fig. 5C), suggesting enhanced tumorigenicity and compromised immunogenicity in dedifferentiated cells. Syngeneic tumors formed by Oct4/Sox2 cells in Balb/c mice exhibited higher level of the proliferation marker Pcna, and c-Myc, Dnmt1, Setdb1, and Ezh2 (Fig. 5D). These proteins promote not only tumorigenicity but also immune evasion of cancer cells. Higher expression of the immune checkpoint Tigit also indicated higher capacity of immune evasion (Fig. 5D). Stronger expression of Afp was also detected, which means higher differentiation potential of the dedifferentiated cells (Fig. 5D). However, there was no significant difference in CD4 or CD8 signal (Fig. 5D), suggesting that response of cancer cells to anti-tumor immunity is not solely mediated by T cell infiltration. By using GFP-labeled cells, we observed that dedifferentiated cells contributed to syngeneic tumor formation similarly to control cells (Fig. 5E), an effect opposite to differentiated cells.

**Fig. 5.**
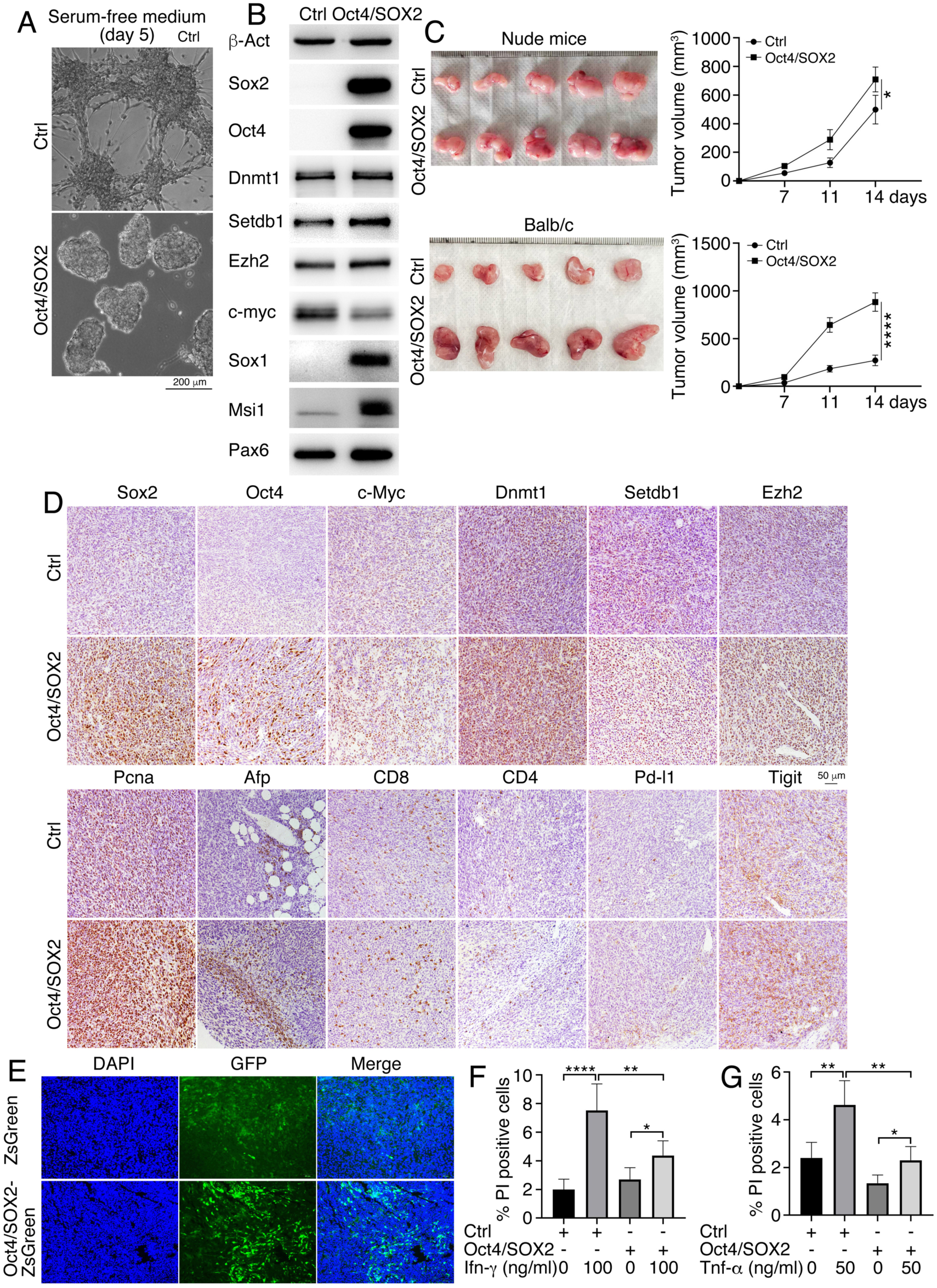
Dedifferentiation changes cellular properties of CT26 cells in the way that is opposite to differentiation. (A) Neurosphere formation of control (Ctrl) cells and cells with simultaneous forced expression of Oct4/SOX2 for 6 days in NSC-specific serum-free culture for 5 days. (B) Dedifferentiation effect after forced expression of Oct4/SOX2 as revealed by WB. (C) Tumor formation in nude and Balb/c mice by control cells and cells with forced expression of Oct4/SOX2. Significance of difference in tumor growth measured with tumor volume was calculated using two-way ANOVA-Bonferroni/Dunn test. Data are shown as mean ± SD. *p < 0.05, ****p < 0.0001. (D) IHC detection of protein or marker expression in sections of tumors formed in Balb/c mice by control and treated cells. Objective magnification: 20 ×. (E) Cryosections showing GFP-labeled cells in tumors formed in Balb/c mice. (F, G) Difference in sensitivity of control and dedifferentiated cells to Ifn-γ (F) or Tnf-α (H) mediated cell death. Significance of difference was calculated using unpaired Student’s *t*-test. Data are shown as mean ± SD. *p < 0.05, **p < 0.01, ****p < 0.0001.

We examined whether dedifferentiation could change the sensitivity to Ifn-γ or Tnf-α mediated cell death. Ifn-γ treatment of either control CT26 or Oct4/SOX2 cells led to increased ratio of dead cells. However, Oct4/SOX2 cells showed lower ratio of dead cells than control cells in response to Ifn-γ treatment (Fig. 5F; Fig. S8C). Similarly, ratio of dead cells was increased in control CT26 or Oct4/SOX2 cells after Tnf-α treatment, but the ratio of dead cells in control CT26 cells was higher than that in Oct4/SOX2 cells (Fig. 5G; Fig. S8C). These data suggest that, contrary to differentiation, dedifferentiation enhances neural stemness and tumorigenicity of CT26 cells, and boosts resistance to cell death mediated by Ifn-γ or Tnf-α.

### MYOD1 suppresses tumorigenesis in the DEN/CCl_4_-induced mouse model of hepatocellular carcinoma

Since differentiation of cancer cells induced by forced expression of lineage-specific differentiation factors or by inhibition of an oncoprotein reprograms their cellular properties and regulatory networks, leading to loss of tumorigenicity and enhanced immunogenicity, we reasoned that differentiation by forced expression of a differentiation factor, for example, MYOD1, could suppress tumorigenesis. We examined whether Myod1 could inhibit DEN/CCl_4_-induced hepatocellular carcinoma (HCC) in mice using adeno-associated virus (AAV) serotype AAV2/8 as the gene delivery vector. The viral vector could effectively express Myod1 when transfected into HEK293 or SK-HEP1 cells (Fig. S9A). Intravenous injection of AAV2/8 showed high transduction efficiency in liver, but not in other organs such as intestine, heart, brain or kidney (Fig. S9B). Mouse model of DEN/CCl_4_-induced hepatocellular carcinoma was made using an established protocol, and AAV particles carrying MYOD1 (AAV-MYOD1) or control AAV (AAV-Ctrl) were administered twice via intravenous injection at indicated time points (Fig. S9C). Tumor formation was observed in both control group that were injected with AAV-Ctrl and the treatment group that were injected with AAV-MYOD1 (Fig. S9D). Nevertheless, liver weight in control group was significantly larger than that in treatment group (Fig. 6A), and the ratio of liver to whole body weight in the control group was also larger than that in treatment group (Fig. 6B). These indexes indicate that the tumor burden in the treatment group was smaller than in control group and MYOD1 suppressed liver tumorigenesis in this model. IHC assays showed that MYOD1 was only detected in the tumors of the treatment group, but not detected in normal liver and tumors of the control group (Fig. 6C). Sox9, Cd133, Msi1, and β-catenin are neural stemness markers and also used as markers for cancer stem cells (CSCs), and Setdb1 expression is enriched in embryonic neural cells. Sox9, β-catenin and Setdb1 are also known as suppressors of anti-tumor immunity (Spranger et al., 2015; Ruiz de Galarreta et al., 2019; Griffin et al., 2021; Zhong et al., 2023). They were absent in normal liver except slight expression of Setdb1 in a part of liver cells, but present strongly in tumor cells of the control group. Tumors with Myod1 expression showed weaker expression of these proteins (Fig. 6C). In normal livers, there was no expression of neuronal markers Map2 and Tubb3, and an endodermal differentiation and HCC marker Afp. Weak express of Acta2, a muscle-specific protein, was present in normal liver. They showed stronger expression in tumors in the control group. In tumors of the treatment group, their expression was decreased (Fig. 6C). Expression of these proteins in normal liver and induced liver cancer reflects the previous proposal that tumorigenesis represents a process of progressive loss of original cell identity and gain of neural stemness, the cellular property that endows cells with not only tumorigenicity but also pluripotent differentiation potential (Cao, 2017; Xu et al., 2021; Cao, 2022; Zhang et al., 2022). The immune checkpoint Pd-l1 was not detected in normal liver, but was present in tumors of the control group. In tumors of the treatment group, it was barely detectable (Fig. 6C). The difference in neural stemness marker and Pd-l1 expression between two groups of tumors was in agreement with that Myod1 suppressed tumorigenesis, and implied that Myod1 enhanced the sensitivity to anti-tumor immunity. Accordingly, CD8+ cells were not detected in normal liver. They were present in tumors of both control and treatment groups, but the signal for CD8 was more strongly detected in tumors with Myod1 expression (Fig. 6C).

**Fig. 6.**
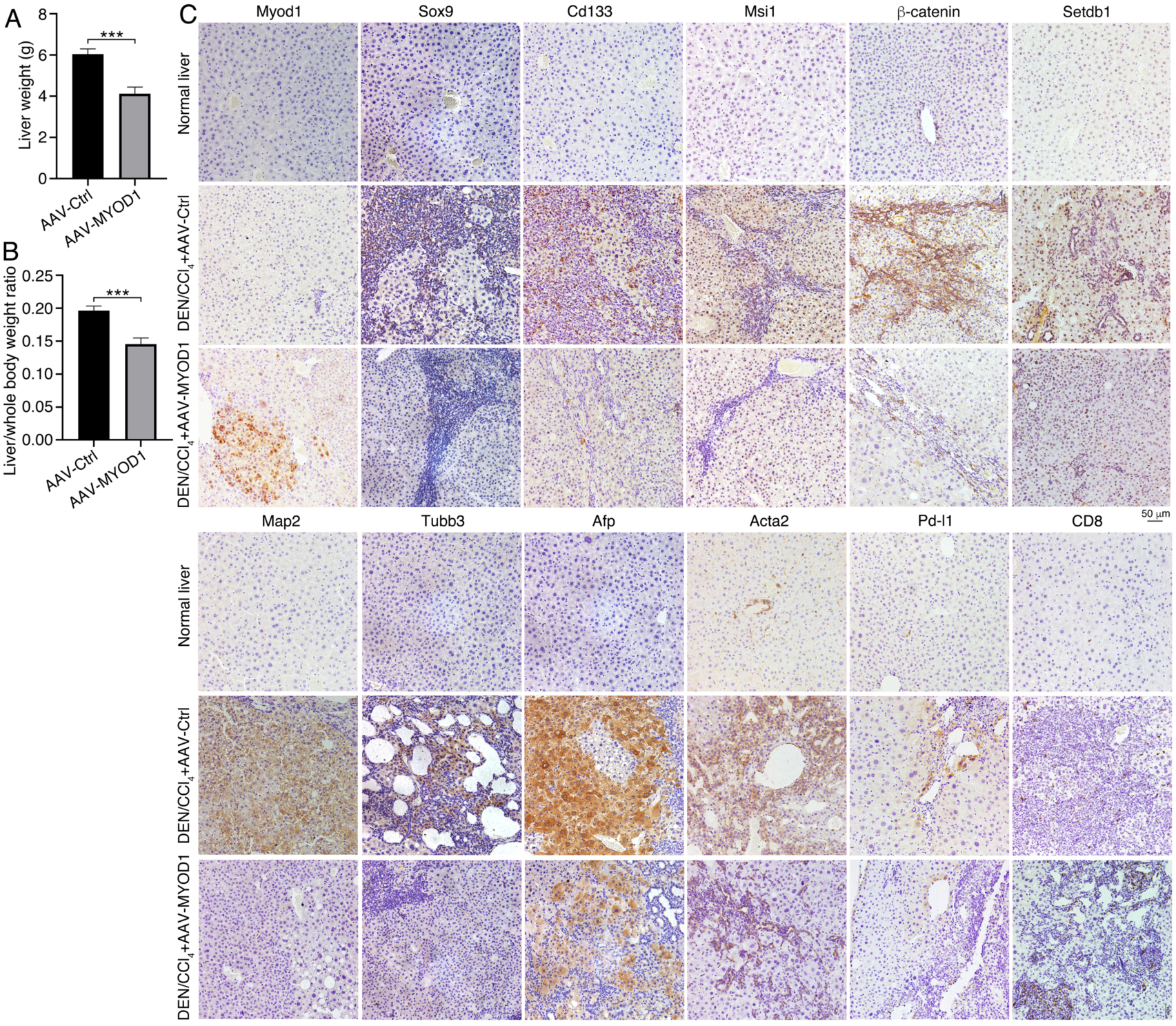
Differentiation factor Myod1 suppresses tumorigenesis in the DEN/CCl_4_-induced mouse model of hepatocellular carcinoma. (A, B) Difference in liver weight (A) or liver/whole body weight ratio (B) between the mice administered with DEN/CCl_4_ and AAV carrying the empty vector (AAV-Ctrl) and the mice administered with DEN/CCl_4_ and AAV carrying MYOD1 (AAV-MYOD1). Significance of difference was calculated using unpaired Student’s *t*-test. Data are shown as mean ± SD. ***p < 0.001. (C) IHC detection of marker expression in normal livers, liver tumors in control group (AAV-Ctrl), and liver tumors in the group that were injected with AAV-MYOD1.

## Discussion

### 1. Pluripotent property of cancer cells dictates that they can be differentiated into various types of cells

CSCs and the property of CSCs, i.e., cancer stemness, are important concepts in cancer biology. However, it was unknown whether CSCs from different types of cancer share a common type of stemness, or CSCs of different cancer types exhibit the tissue stemness of their respective tissues of cancer origin, or CSCs are not comparable with any known stem/progenitor cell types (Cao, 2022). In history, the earliest characterized property of cancer stemness was the pluripotency of the cells of teratocarcinoma, a type of cancer found in many different tissues. Now pluripotency has become a keyword in developmental/stem cell biology, it faded into oblivion in mainstream cancer research. Our studies showed that the core property of cancer (tumorigenic) cells is the evolutionarily predestined neural stemness, which represents the general stemness and determines both pluripotent differentiation potential and tumorigenicity (Cao, 2017; Cao, 2022; Cao, 2023; Chen et al., 2021; Xu et al., 2021; Zhang et al., 2017; Zhang et al., 2022). This means that, like teratocarcinoma or pluripotent embryonic cells, cancer cells can be differentiated into different cell types. Our studies have demonstrated that different types of cancer cells can be induced to differentiate along different lineages, either by forced expression of lineage specific specification factors or by blocking endogenous oncoproteins (Zhang et al., 2017; Lei et al., 2019; Zhang et al., 2022; Chen et al., 2021; present study). Immune cells are the key player in anti-cancer immunity. Differentiation of cancer cells into cells resembling immune cells was predicted based on cancer cell pluripotency (Cao, 2023). Indeed, in vitro and in vivo experimental data verified that cancer cells can be reprogrammed into antigen-presenting cells by lineage-specific specification factors (Ascic et al., 2024; Linde et al., 2023; Zimmermannova et al., 2023). A latest research showed induced differentiation of colorectal cancer cells into normal-like enterocytes, thereby suppressing malignancy (Gong et al., 2025). Taken together, our and other studies have confirmed the pluripotent property of cancer cells. In principle, they can differentiate into any types of cells, following the rules of cell/tissue differentiation during embryonic development.

### 2. Cellular properties of cancer cells, including tumorigenicity and immunogenicity, are coupled together and associated with the differentiation status

Cancer cells display different cellular properties, such as tumorigenicity, pluripotency, fast proliferation, metastasis, evasion of death and anti-cancer immunity, dysregulated metabolism and epigenetics, therapy resistance. It is common to see that a same oncofactor regulates different properties of cancer cells. For example, EZH2 plays oncogenic role in different types of cancers, promotes cancer cell stemness (Balinth et al., 2022), proliferation (Bryant et al., 2007), metastasis (Zingg et al., 2015), chemoresistance (Ougolkov et al., 2008; Crea et al., 2012), metabolism (Ahmad et al., 2017), etc. Similarly, numerous studies on c-Myc revealed that it regulates almost all properties of cancer cells (Dhanasekaran et al., 2022; Fatma et al., 2022; Llombart and Mansour, 2022). Cancer immunity and metabolism are prevailing research fields in present cancer biology. Typical oncoproteins are also major regulators of cancer cell metabolism and immunogenicity, e.g., EZH2 (Kim et al., 2020; Nylund et al., 2021; Zhou et al., 2020), C-MYC (Miller et al., 2012; Zimmerli et al., 2022), KRAS (Kerk et al., 2021; Watterson and Coelho, 2023; Lasse-Opsahl et al., 2025). A same factor being able to regulate different properties of cancer cells suggests that these properties are not independent from each other. All these cancer promoting factors are components of embryonic neural regulatory network, which endows cancer cells with neural stemness (Cao, 2017; Cao, 2022; Xu et al., 2021; Zhang et al., 2017; Zhang et al., 2022). This means that different cellular properties of cancer cells are ultimately determined and coupled together by neural stemness and its corresponding regulatory networks. Disrupting one cellular property of cancer cells will inevitably affect one or more, if not all, other properties, as shown in present and many other studies.

Neural stemness being the core property of cancer cells means that genes responsible for differentiation, cellular properties and functions of differentiated cells are generally silenced in cancer cells by cancer promoting genes. Disruption of neural regulatory network in cancer cells via inhibition of one or more endogenous cancer promoting factors, as in the strategy of targeted therapies, or forced expression of one or more differentiation factors leads to stimulation of differentiation genes, leading to the loss of neural stemness and acquirement of the properties of differentiated cells that are not tumorigenic anymore (Zhang et al., 2017; Krah et al., 2019; Lei et al., 2019; Chen et al., 2021; Yang et al., 2021; Zhang et al., 2022; Linde et al., 2023; Zimmermannova et al., 2023; Ascic et al., 2024; He et al., 2025; present study). Immunogenicity of cells relies on the expression of immune related genes. The present study identifies a general tendency of upregulation of immune related genes and hence enhanced immunogenicity in cancer cells after differentiation. Vice versa, dedifferentiation enhances neural stemness and tumorigenicity, but suppresses immune related genes and immunogenicity. Therefore, tumorigenicity and immunogenicity are inversely correlated, and both are determined by the differentiation status of cancer cells. This general rule is also reflected by normal NSCs and differentiated cells because the former are tumorigenic and immune-privileged and the latter are just the opposite. The studies on differentiation of cancer cells into antigen-presenting cells analyzed only their ability to promote anti-cancer immunity and underlying mechanisms in the context of immunity, but leaving tumorigenicity untested (Ascic et al., 2024; Linde et al., 2023). However, it can be predicted that cancer cells after differentiation into immune cells will inevitably lose their tumorigenicity, as shown by another study (Zimmermannova et al., 2023). Loss of tumorigenicity in cancer cells means clearly a tumor suppression effect. Interpretation of this effect by focusing merely on specific pathways of anti-cancer immunity suggests that response of cancer cells to anti-cancer immunity is the most important property, but tumorigenicity is not. However, differentiated cancer cells lost tumorigenicity even in an environment of severely impaired immunity, such as in mice lacking T, NK and macrophage cells (Zimmermannova et al., 2023). This effect should not be interpreted purely as the consequence of anti-tumor immunity.

### 3. Enhanced immunogenicity by differentiation

A primary research focus on cancer immunotherapy is to find out ways to boost the sensitivity of cancer cells to anti-cancer immunity. Increasing data confirmed the critical role of major cancer promoting factors in promoting immune evasion and immunotherapy resistance of cancer cells via pairwise molecular regulatory mechanisms. For instances, C-MYC blunts immune cell invasion in triple-negative breast cancer via suppression of STING-IFN signaling in a tumor cell-intrinsic fashion (Zimmerli et al., 2022); Inhibition of CDK4/6 de-represses NFAT family proteins, thereby enhancing T cell activation (Deng et al, 2018); β-catenin prevents anti-cancer immunity via repressing *CCL4* transcription (Spranger et al., 2015); Inhibition of KRAS(G12D) induces FAS expression in cancer cells and facilitates CD8+ T cell-mediated death (Mahadevan et al., 2023); Inhibition of EZH2 upregulates MHC class I expression and increases antigen-specific CD8+ T-cell proliferation, IFNγ production, and tumor cell cytotoxicity (Zhou et al., 2020); HDAC1 controls anti-cancer immunity by suppressing type I dendritic cell maturation (De Sá Fernandes et al., 2024). Due to the complexity of cancer and immunity, these pairwise regulatory mechanisms might be countless. It is critical to know whether there is a general rule governing immunogenicity of cancer cells. Although cancer stemness is considered as a key factor driving immune evasion and immunotherapy resistance of cancer cells (Galassi et al., 2021; Agudo ands Miao, 2024), the intrinsic link is still difficult to establish without clarification of the essence of cancer stemness. Neural stemness as the core property of cancer cells facilitates characterization of their cellular properties, including immunogenicity. According to principles of developmental biology, different types of cells are determined by their respective cell type specific regulatory networks, exhibit distinct cellular properties and execute different functions. There exist general antagonisms of expression of genes promoting cell fate specification/differentiation, maintaining cell identity and function between different type cells, thereby guaranteeing tissue or cell identity. Therefore, neural stemness is determined by genes specifically expressed in or enriched in neural stem cells/embryonic neural cells, which are generally repressed or silenced in non-neural cells. Vice versa, genes specifying non-neural cells or maintaining their identities/functions, including immune related genes, are not highly expressed in embryonic neural cells. Such a correlation is confirmed by the analysis on the transcriptomes of neural stem cells, embryonic stem cells, muscle, fat, spleen and thymus cells. Interestingly, immune related genes, particularly those enhancing immunogenicity, including IFN-γ response genes and genes involved in antigen processing and presentation, are not or weakly expressed in neural stem or embryonic stem cells, but they are highly expressed in the organs of the immune system and in other non-neural cells, e.g., muscle and fat cells. This difference in immune related gene expression is in agreement with that neural stem cells and embryonic stem cells, whose default fate is neural stem cells, are immune privileged, but other types of cells are not (Fändrich et al., 2002; Hori et al., 2003; Drukker et al., 2006; Magliocca et al., 2006; Itakura et al., 2017; Ozaki et al., 2017). Therefore, neural stemness endows cancer cells with not only tumorigenicity, pluripotency and other malignant features, but also immune privilege. Differentiation of cancer cells, either by forced expression of a lineage-specific differentiation factor or by inhibition of endogenous cancer promoting factor, led to reprogramming the regulatory networks of cancer cells into those of differentiated cells, and reprogramming the cellular properties of cancer cells into the properties of differentiated cells with reduced neural stemness and tumorigenicity. Moreover, differentiation effect caused a general tendency of decreased expression of cancer promoting genes, which promotes immune evasion, and increased expression of immune related genes including those enhancing immunogenicity, thereby enhancing the sensitivity of cancer cells to IFN-γ or TNF-α mediated cell death. As a result, a majority of cancer cells after differentiation showed failure in contribution to tumor formation in syngeneic and allogeneic transplantation assays. By contrast, dedifferentiation causes upregulation of cancer promoting genes and neural stemness genes, and change transcriptomic profile more characteristic of neural stem cells. Dedifferentiated cells show enhancement of neural stemness, malignant features and tumorigenicity. Meanwhile, immune related genes are downregulated, leading to cells with less sentivity to IFN-γ and TNF-α mediated cell death. Therefore, tumorigenicity and immunogenicity are both determined by differentiation status of cancer cells.

The molecular mechanisms of immune evasion regulated by cancer promoting factors or enhanced immunogenicity in response to inhibition of cancer promoting factors actually reflect the intrinsic link between neural stemness, differentiation status and immunogenicity of cancer cells. Most (if not all) cancer promoting genes, such as *MYC*, *β-catenin*, *CDK4/6*, *KRAS*, *EZH2*, and *HDAC1*, are enriched in neural stem/embryonic neural cells (Cao, 2022) and play roles in specifying neural stem/precursor cells and/or maintaining neural stemness. On one hand, they promote immune evasion by promoting the cancer regulatory network, i.e., the embryonic neural network, and on the other, they are involved in repressing non-neural genes, including immune related genes in neural stem cells and cancer cells. Blocking one or more components in the regulatory network in cancer cells will lead to a differentiation effect and de-repression of repressed genes, including immune related genes, thereby enhancing immunogenicity of cancer cells. It is appropriate to say that upregulation of immune related genes, e.g., MHC genes, and hence enhancement of anti-cancer immune response via inhibition of oncofactors is per se the consequence of differentiation. Inhibition of cancer promoting factors in cancer cells already causes suppression of tumorigenicity, enhanced sensitivity to apoptosis or therapy resistance, as revealed by numerous studies, but not merely an effect of enhanced immunogenicity. However, most commonly seen individual studies focused on a particular cellular property of cancer cells under the regulation of a particular molecule or pairwise regulatory axis. Our studies have shown that the major cellular properties of cancer cells, e.g., tumorigenicity, pluripotency, and immunogenicity, are coupled together by neural stemness and its corresponding regulatory networks. Cells with neural stemness express high level of neural stemness/embryonic neural genes but low level of pro-differentiation genes and genes specifying/maintaining the functions of differentiated cells, including immune related genes, thereby conferring these cells with tumorigenicity, pluripotency and immune privilege. Upon differentiation, cells will lose neural stemness with reduced expression of neural stemness/embryonic neural genes and elevated expression of pro-differentiation genes and genes specifying/maintaining the functions of differentiated cells, including immune related genes. The resulting cells thus exhibit reduced tumorigenicity and pluripotency and enhanced immunogenicity. These results reinforce neural stemness as the core property of cancer cells, which determines other basic properties of cancer cells, including pluripotency, tumorigenicity, and immunogenicity. The general relation between neural stemness, differentiation status and cancer cell properties, or the **cellular mechanism** underlying tumorigenicity and immunogenicity of cancer cells, was summarized in Figure 7 (Fig. 7).

**Fig. 7.**
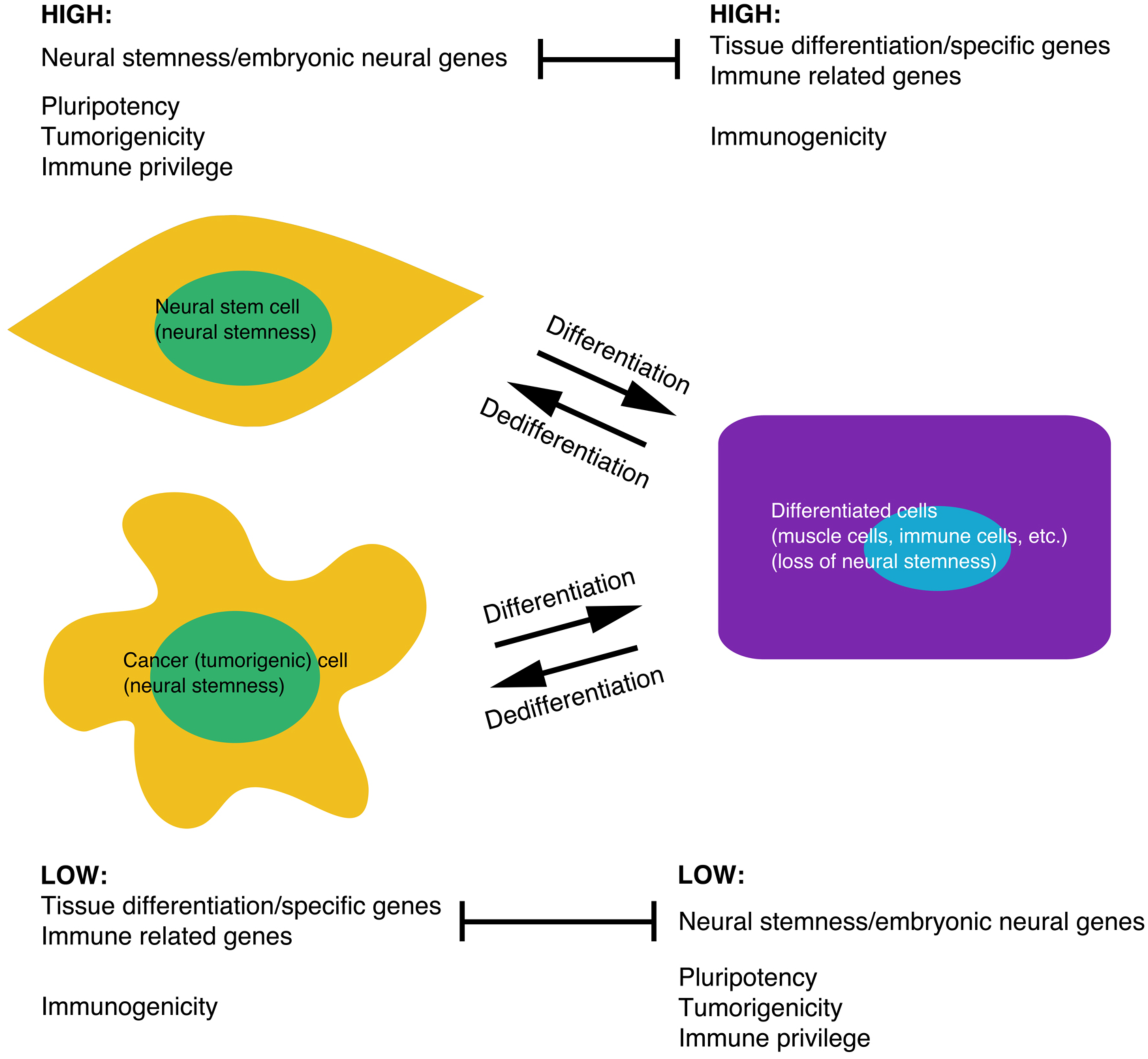
A model depicting the general relationship between differentiation status and cellular properties, particularly tumorigenicity and immunogenicity, of cancer cells. Stemness of neural stem cells, i.e., neural stemness, is defined by neural stem or embryonic neural genes and represents general stemness, which confers cells with pluripotent differentiation and tumorigenic potentials (Cao, 2022; Zhang et al., 2022). Meanwhile, immune related genes are not or only weakly expressed in neural stem cells, conferring cells with immune privilege. In differentiated cells, neural stemness is lost and neural stem or embryonic neural genes are repressed, but tissue differentiation or tissue specific genes are stimulated, conferring cell-specific properties or functions to cells. Accordingly, differentiated cells lose pluripotency and tumorigenicity. Meanwhile, immune related genes are also stimulated, leading to a loss of immune privilege and acquirement of immunogenicity in differentiated cells. The relationship between neural stemness/embryonic neural genes and immune related genes reflects the general rule that genes specifying the properties and functions of different types of cells should antagonize mutually to guarantee the identity and function of a type of cells and establish boundaries between different tissues and organs. The core property of cancer (tumorigenic) cells is neural stemness. They are pluripotent, tumorigenic and immune privileged cells. The mutually exclusive transcription of neural stemness/embryonic neural genes and tissue differentiation/specific genes and immune related genes, and contrasting cellular properties defined by these genes, including differentiation potential, tumorigenicity, and immunogenicity, are also manifested in undifferentiated and differentiated cancer cells. Differentiation causes the loss of neural stemness, differentiation potential, tumorigenicity and immune privilege due to downregulation of neural stemness/embryonic neural genes and upregulation of differentiation/cell-specific genes and immune related genes, and dedifferentiation causes an opposite effect. This general rule emphasizes that cancer should be understood by understanding neural stemness and underlying developmental principles controlling it, and that targeting neural stemness via induced differentiation should be efficient strategy of cancer therapy because of suppression of tumorigenicity and meanwhile enhancement of immunogenicity of cancer cells.

### 4. Differentiation therapy of cancer via targeting neural stemness

Targeting neural stemness by induced differentiation of cancer cells will reprogram their cellular properties and regulatory networks, suggesting a promising strategy for cancer therapy. Differentiation therapy is not a new concept. It was already suggested 50 years ago (Pierce and Wallace, 1971). Treatment of acute promyelocytic leukemia with all-trans retinoic acid might be the best-known case of differentiation therapy (de Thé, 2018). The major obstacle might be the inappropriate understanding of the key property of cancer cells. The core property of cancer cells being neural stemness, which endows cells with pluripotency, provides a general framework for differentiation therapy of different types of cancers: cancer cells can be induced to differentiate by differentiation factors, particularly those driving embryonic tissue differentiation. A series of studies in the 1970-80s demonstrated differentiation of cancer cells into benign cells within embryonic environment (Cao, 2023). In recent years, cancer cells were shown to differentiate into different types of cells induced by lineage differentiation factors, inhibition of cell-intrinsic cancer promoting factors or by chemical molecules (Zhang et al., 2017; Krah et al., 2019; Lei et al., 2019; Chen et al., 2021; Yang et al., 2021; Zhang et al., 2022; Linde et al., 2023; Zimmermannova et al., 2023; Ascic et al., 2024; Gong et al., 2025; He et al., 2025). Targeted therapies via inhibition of endogenous cancer promoting factors are in essence differentiation therapies. The efficiency of cancer therapy via inhibition of endogenous cancer promoting factors relies on the uniformity of presence of a target in cancer cells and the degree of differentiation of cancer cells in response to target inhibition. In addition, cancer with inhibition of a target gene/protein might be more prone to signal feedback loops in cancer regulatory networks, which will ultimately lead to therapy resistance. By contrast, induced differentiation driven by embryonic non-neural differentiation factors does not depend on a specific target protein, and can achieve more efficient differentiation effect, reprogram overall cellular properties and regulatory networks of cancer cells, and might avoid or alleviate therapy resistance effect caused by complex signal feedback loops. We showed previously that a few embryonic non-neural lineage differentiation factors, GATA3, HNF4A, HHEX, and FOXA3, suppress tumorigenicity via inhibition of neural stemness of cancer cells, and suppress tumorigenesis in a mouse model of colorectal cancer (Yang et al., 2021). The present study revealed that tumorigenicity and immune privilege of cancer cells is coupled together by neural stemness, and differentiation status of cancer cells dictates tumorigenicity and immunogenicity. Moreover, the muscle differentiation factor Myod1 suppresses tumorigenesis in a mouse model of hepatocellular carcinoma. In summary, neural stemness and its regulatory network determines basic cellular properties of cancer cells, including tumorigenicity, pluripotency and immune privilege by repressing differentiation genes, including immune related genes. Induced differentiation of cancer cells suppresses cancer promoting genes and re-activates differentiation genes, leading to loss of neural stemness and hence tumorigenicity, pluripotency and immune privilege, and gain of the properties of differentiated cells, including enhanced immunogenicity. Regulation of immunogenicity or tumorigenicity and the underlying molecular mechanisms is complex, and specific molecular mechanisms represented by particular genes/proteins are specific details of the complicated scenario. We identified the **cellular mechanism** underlying the properties of cancer cells, particularly immunogenicity and tumorigenicity. This suggests to understand cancer by understanding neural stemness and the principle of embryonic cell/tissue differentiation, and suggests to develop novel therapies for cancer treatment based on the principle of differentiation.

## Materials and methods

### Cell culture

HEK293T, HCT116, SK-HEP1, and B16F10 cells were cultured in Dulbecco’s modified eagle medium (DMEM. Thermo Fisher Scientific, #11965-092), and CT26 cells were in RPMI-1640 medium. All media were supplemented with 10% fetal bovine serum (FBS. Gibco, #10099141) and 50 U/ml penicillin/50 µg/ml streptomycin. Mouse embryonic stem cells (mESCs) were cultured in DMEM medium, supplemented with 15% FBS, 1 ng/ml human LIF (Cell Signaling Technology, #8911), 2 mM L-glutamine (Thermo Fisher Scientific, #25030164), 100 μM β-mercaptoethanol, 1× MEM non-essential amino acids, and 50 U/ml penicillin/50 µg/ml streptomycin. For culture of mESCs, petri dishes were coated with 0.1% gelatin. All cells were cultured at 37°C with 5% CO_2_. HEK293T (Cat. No.: #SCSP-502), HCT116 (Cat. No.: #TCHu 99), SK-HEP1 (Cat. No.: #TCHu 109), B16F10 (Cat. No.: #SCSP-5233), CT26 (Cat. No.: #TCM 37) and mESCs (Cat. No.: #SCSP-218) were purchased from the Cellbank of Chinese Academy of Sciences (Shanghai, China). Cancer cell lines were authenticated with short tandem repeat profiling. Cells with fewer than eight passages were used for experiments.

Cells were also cultured in a defined serum-free medium Ndiff^®^227 (CellArtis, #Y40002) used for derivation of primitive neural stem cells (primNSCs) from mESCs and for the test of neurosphere-like structure formation by cancer cells.

### Extraction of mouse tissue cells

Muscle, fat, spleen and thymus tissues were obtained from B6 male mice at 4 weeks of age. Skeletal muscle tissue was removed from legs and washed with pre-cooled PBS (Phosphate buffered saline) under sterile conditions. Muscle was then digested with 0.2% type I collagenase at 37°C with 5% CO_2_ for 50 minutes to obtain isolated muscle fibers. After neutralizing digestion with serum-containing DMEM and washing with PBS, muscle fibers were collected. Subcutaneous fat tissue was removed from the groin and washed three times with PBS. Fat tissue was cut into small pieces, and digested with 0.1% type I collagenase in a water bath at 37°C for 40 minutes. After termination of digestion, the mixture was filtered through a 100-mesh filter, and centrifuged at 140×g for 5 min. The upper layer of fat cells was collected and washed with PBS. To obtain spleen cells, spleen tissue was pulverized, filtered, centrifuged at 500×g for 5 minutes, and the supernatant was discarded. 500 μl red blood cell lysis buffer was added to the precipitated cells, treated at room temperature for 10 minutes to remove red blood cells. Spleen cells were then collected by centrifugation and PBS washing. Muscle fibers, fat cells, spleen cells and thymus tissue were subjected to RNA-sequencing.

### Plasmid construction, virus packaging, and cell transduction

Validated MISSION^®^ short-hairpin RNA (shRNA) (Sigma-Aldrich) TRCN0000092973 was used for knockdown of mouse Setdb1, and TRCN0000276105 was used for knockdown of human SETDB1. shRNAs were subcloned to the lentiviral vector pLKO.1 and designated as shSetdb1 and shSETDB1, respectively.

For stable expression of human MYOD1 or PPARG or mouse Myod1 or Pparg, the coding regions of MYOD1/Myod1 or PPARG/Pparg were subcloned to lentiviral vectors pLVX-IRES-Puro or pLVX-IRES-ZsGreen. For double expression of Oct4 and SOX2 in cells, coding regions of Oct4 and SOX2 were linked together by P2A sequence and subcloned to pLVX-IRES-Puro or pLVX-IRES-ZsGreen.

Lentivirus production and cell infection were performed according to polyethylenimine (PEI) transfection protocol. HEK293T cells were co-transfected with packaging plasmids and shRNA or expression constructs in the presence of PEI. 48 hours after transfection, lentiviral supernatant was filtered through 0.45 μm filters and centrifuged at 4°C to concentrate lentiviral particles, which were used for transducing cells. Viral transduction of cells was performed in the presence of polybrene at a final concentration of 10 µg/ml. Cells after transduction for 48 hours were selected with puromycin at different concentrations (1 μg/ml for HCT116, 2 μg/ml for SK-HEP1 and B16F10, and 10 μg/ml for CT26) in culture for three days when a puromycin selection vector was used. Cells were cultured further for observation of phenotypic change or for additional assays. As a control for knockdown or protein expression assays, virus production with empty vector and cell transduction were performed in parallel.

For testing tumor suppression activity of MYOD1 in mouse model of hepatocellular carcinoma, the coding region of MYOD1 was subcloned to adeno-associated virus (AAV) vector pAAV-CBh-EGFP-WPRE to generate the construct pAAV-CBh-3xFLAG-MYOD1-P2A-EGFP-WPRE. Construction of AAV construct and AAV packaging and tittering was performed by OBiO Technology (Shanghai, China).

### Oil red staining

Oil red O staining was performed to detected adipocyte differentiation of cancer cells induced by PPARG/Pparg expression. Control and treated cells were cultured on coverslips in 6-well plates for a desired period. Cells were then washed with PBS for three times, followed by fixation with 4% PFA for 30 min, and again three times of wash with PBS. Afterwards, cells were rinsed with 60% isopropyl alcohol for one minute, and stained with freshly prepared oil-red O working solution for 30 min when lipid staining could be observed.

### Immunoblotting

Whole cell lysates were prepared for detecting protein expression using conventional SDS-PAGE and immunoblotting. Protein bands were revealed with a Western blotting substrate (Tanon, #180-501). Primary antibodies were: β-ACT (ABclonalCell Signaling Technology, #4970. 1:10,000), DNMT1 (ABclonal, A16729, 1:1,000), EZH2 (Cell Signaling Technology, #5246. 1:2,000), FABP4 (ABclonal, A0232. 1:1,000),, MAP2 (Abcam, #ab183830. 1:1,000), MEF2C (Cell Signaling Technology, #5030. 1:1,000), MYOD1 (Novus Biologicals, NBP1-54153. 1:1,000), MYOG (Abcam, # ab124800. 1:1,000), MSI1 (ABclonal, A9122. 1:1,000), C-MYC (Abcam, #ab42072. 1:1,000), NEUN (Cell Signaling Technology, #12943. 1:1,000), Oct4 (Cell Signaling Technology, #83932. 1:1,000), PAX6 (Abcam, #ab195045. 1:1,000), PCNA (Cell Signaling Technology, #13110. 1:2,000), PPARG (Cell Signaling Technology, #2435. 1:1,000), SETDB1 (Cell Signaling Technology, #2196. 1:1,000), SOX1 (Abcam, #ab109290. 1:1,000), SOX2 (Abcam, #ab92494. 1:1,000), SYN1 (Cell Signaling Technology, #5297. 1:1,000), VGLUT1 (Cell Signaling Technology, #12331. 1:1,000).

### Cell migration/invasion assay, and soft agar colony formation assay

These assays were performed essentially as described (Zhang et al., 2022). Cell migration assay was carried out using 24-well transwell plates with inserts of 8-µm pore size (Biofil, #TCS003024). The upper compartment of a well contained 1×10^5^ control or treated cells that were suspended in 200 µl of culture medium without FBS. The lower compartment contained 500 µl of culture medium supplemented with 10% FBS. Plates were incubated at 37°C for desired period as indicated in the text. Afterwards, cells were washed with PBS, fixed with 37% formaldehyde, and stained with 0.5% crystal violet for 5 min. Cells that didn’t migrate were removed. Migrated cells were washed with PBS, and observed with microscopy.

For cell invasion assay, each 5×10^5^ control or treated cells were added to 80 µl of Matrigel (Corning, #354234), then diluted in PBS at a ratio of 1:8, and evenly distributed onto a 24-well transwell insert. They were incubated at 37°C for a desired period, as indicated in the text. Afterwards, cells were processed in the same way as in the migration assay.

For colony formation assay, soft agar was made of two layers of low melting agarose (Sangon Biotech, #A600015). The top layer was 0.35% of agarose in complete culture medium, and the bottom layer was 0.7% of agarose. In each well of a 6-well culture plate, 2,000 control or treated cells were plated on the top layer of agar, and cultured at 37°C for a desired period as indicated in the text. Experiments were repeated three times. Colonies larger than 50 µm or 100 µm in diameter, depending on the type of cells, were counted for analysis using unpaired Student’s *t*-test.

### Cancer cell transplantation

Mouse use in the study was approved by the Institutional Animal Care and Use Committee (IACUC) at the Model Animal Research Center of Medical School, Nanjing University, and experiments with mice were performed in accordance with the guidelines of IACUC. Immunodeficient nude Foxn1^nu^ male mice of 5-6 weeks old, C57BL/6J, and Balb/c mice of 5-6 weeks old were purchased from the National Resource Center for Mutant Mice (Nanjing, China) and maintained in specific-pathogen-free facility. For xenograft tumor assay, control or treated cancer cells were suspended in 100 μl of PBS and injected subcutaneously into the dorsal flank of mice. For syngeneic transplantation, control or treated B16F10 cells were injected subcutaneously into the dorsal flank of C57BL/6J mice, or control or treated CT26 cells were injected subcutaneously into the dorsal flank of Balb/c mice. For allogeneic transplantation, control or treated B16F10 cells were injected subcutaneously into the dorsal flank of Balb/c mice, or control or treated CT26 cells were injected subcutaneously into the dorsal flank of C57BL/6J mice. Tumor formation was observed and tumor size was measured periodically before mice were sacrificed. After sacrifice of mice, tumors were excised. Tumor volume was calculated with the formula: length×width^2^/2. Significance of difference in tumor volume between control and treated groups was calculated with two-way ANOVA-Bonferroni/Dunn test. Control or treated HCT116 cells were injected at a dose of 3×10^6^ cells per mouse, SK-HEP1 at a dose of 3×10^6^, B16F10 at 5×10^5^, and CT26 injected at a dose of 5×10^5^ per mouse.

### Immunohistochemistry (IHC)

IHC was used to detect expression of proteins in paraffin sections of tumors according to conventional method. Briefly, paraffin in sections was removed by washing in xylene for 10 min first and then 5 min. Sections were rehydrated with sequential washes in 100%, 95%, 85%, 70%, 50% ethanol, and dH2O. Antigen retrieval was performed by treating slides with 0.01 M sodium citrate solution at 95-100°C for 20 min. Slides were rinsed with PBS after cooling to room temperature. Endogenous peroxidase activity was blocked with 3% H_2_O_2_ in methanol at room temperature for 15 min, followed by washing slides with PBS three times. Sections were blocked with 5% BSA in PBS for 1 hour at room temperature, then incubated with primary antibody at 4°C overnight, and washed with PBS. Biotin-conjugated goat anti-rabbit secondary antibody (Sangon Biotech, #D110066. 1:500) was added to sections. After incubation at room temperature for 1 hour, a DAB substrate (Sangon Biotech, #E670033) was added. Sections were incubated at room temperature for a desired period (3-15 min) for signal visualization. Cell nuclei were counterstained with hematoxylin. Primary antibodies were: ACTA2 (Abclonal, #A11111. 1:250), AFP (Cell Signaling Technology, #4448. 1:250), β-Catenin (non-phosphorylated) (Cell Signaling Technology, #8814. 1:800), CD133 (Novus Biologicals, NB120-16518. 1:200), CD8 (Abcam, #ab217344. 1:200), C-MYC (Abcam, ab32072. 1:200), DNMT1 (ABclonal, A16729, 1:200), KI67 (Cell Signaling Technology, #9129. 1:200), MAP2 (Abcam, #ab183830. 1:200), MYOD1 (Novus Biologicals,NBP1-54153. 1:200), MSI1 (ABclonal, A9122. 1:200), PCNA (Cell Signaling Technology, #13110. 1:200), PD-L1 (Cell Signaling Technology, #13684, #64988. 1:200), PPARG (Cell Signaling Technology, #2435. 1:200), SETDB1 (Cell Signaling Technology, #2196. 1:200), SOX1 (Abcam, #ab109290. 1:200), SOX9 (Cell Signaling Technology, #82630. 1:200), TIGIT (Cell Signaling Technology, #99567; Abcam, #ab30073. 1:200), TUBB3 (Cell Signaling Technology, #4466. 1:200).

### Cryosection

Tumors were fixed in 4%PFA, dehydrated in 30% sucrose, embedded in Tissue-Tek OCT (Sakura, #4583), and frozen in liquid nitrogen. Cryosections were made with a cryostat microtome (Leica). For fluorescence detection, sections were placed at room temperature for 30 min and then washed with PBS. Cell nuclei were stained with DAPI. Fluorescence was observed under a fluorescence microscope (Olympus BX53).

### Transcriptome profiling

Transcriptomes of control HCT116, SKHEP1, B16F10, and CT26 cells, and cells with forced expression of MYOD1, PPARG, or Oct4/SOX2, or knockdown of SETDB1 (as described in the text) were analyzed with RNA sequencing and their differences were compared. Transcriptomes of mouse ESCs, primitive NSCs derived from mESCs, and muscle, fat, thymus and spleen cells that were isolated from mouse were also analyzed with RNA sequencing and compared.

Total RNA was prepared with TRIzol. Total amounts and integrity of RNA were assessed using the RNA Nano 6000 Assay Kit of the Bioanalyzer 2100 system (Agilent Technologies, CA, USA). mRNA was purified from total RNA by using poly(T) oligo-attached magnetic beads. After fragmentation of mRNA, first strand cDNA was synthesized using random hexamer primer and M-MuLV reverse transcriptase. Subsequently, RNA was degraded using RNaseH, and second strand cDNA synthesis was performed using DNA Polymerase I and dNTP. cDNA overhangs were blunted, 3’ ends of DNA fragments were adenylated. Adaptors with hairpin loop structure were ligated to prepare for hybridization. The library fragments were purified with AMPure XP system (Beckman Coulter, Beverly, USA) to select cDNA fragments of preferentially 370-420 bp in length. After qualification, cDNA library was subjected to sequencing by Illumina NovaSeq 6000. The end reading of 150bp pairing was generated. The image data measured by the sequencer were converted into sequence data (reads) by CASAVA base recognition. Raw data (raw reads) of fastq format were firstly processed through in-house perl scripts, and mapped against mouse or human reference genomes using Hisat2 (v2.0.5) software. The mapped reads of each sample were assembled by StringTie (v1.3.3b) (Mihaela Pertea.et al. 2015) in a reference-based approach. The featureCounts v1.5.0-p3 was used to count the reads numbers mapped to each gene. FPKM of each gene was calculated based on the length of the gene and reads count mapped to this gene. Differential expression analysis of two conditions was performed using the edgeR package (3.24.3). P values were adjusted using the Benjamini-Hochberg method. Padj ≤ 0.05 and |log2(fold change)| ≥ 1 were set as the threshold for significantly differential expression.

Gene Ontology (GO) and KEGG pathway enrichment analysis of differentially expressed genes was performed with the clusterProfiler R package (3.8.1). Gene Set Enrichment Analysis (GSEA) was performed by using the local version of the GSEA analysis tool (http://www.broadinstitute.org/gsea/index.jsp) to determine if a predefined Gene Set can show a significant consistent difference between two biological states. The genes were ranked according to the degree of differential expression in the two samples, and then the predefined Gene Set were tested to see if they were enriched at the top or bottom of the list. Sequencing, signal processing and data analyses were performed by Novogene Co., Ltd. (Beijing, China). RNA-seq data were deposited to Gene Expression Omnibus (GEO) under accession numbers GSE295152, GSE295153, GSE295154, GSE295155, GSE295156, GSE295157, GSE295158, GSE295159, and GSE290839).

Differentially expressed IFN-γ response genes and antigen processing and presentation genes were identified according to the gene sets of “HALLMARK_INTERFERON_GAMMA_RESPONSE” and “GOBP_ANTIGEN_PROCESSING_AND_PRESENTATION” in the GSEA database (https://www.gsea-msigdb.org/gsea/msigdb). Genes involved in TNF-α mediated cell death were from van Loo and Bertrand, 2023 (van Loo and Bertrand, 2023).

### DEN/CCl_4_-induced model of hepatocellular carcinoma

Male B6 mice at 2 weeks were separated into three groups. One group of mice was not treated and used as negative control. Other two groups were injected intraperitoneally with DEN (diethylnitrosamine) at a dose of 25 mg/kg body weight per mouse. Since 4 weeks of age, intraperitoneal injection of CCl_4_ (5 μl/g. CCl_4_:olive oil=1:4) was made twice a week and made for 16 consecutive weeks. First AAV injection was performed at the age of 16 weeks. One group of mice was injected intravenously with control AAV (AAV-Ctrl) at a dose of 2 × 10^11^ v.g./100 μl per mouse, the other group was injected intravenously with AAV for MYOD1 expression (AAV-MYOD1) at the same dose. AAV injection was repeated once when mice reached the age of 20 weeks. At 30 weeks of age, mice were sacrificed and liver tissues were dissected, measured, and then fixed for further analyses. Difference in liver weight or in the ratio of liver/whole body weight was analyzed with unpaired Student’s *t*-test.

### IFN-γ and TNF-α treatment of cells

Control B16F10 or CT26 cells and cells with induced differentiation/dedifferentiation were seeded into 48-well plates, and treated with IFN-γ (PepRotech, #315-05) (20 ng/ml for B16F10, and 100 ng/ml for CT26) or TNF-α (PepRotech, #AF-315-01A) (100 ng/ml for B16F10, and 50 ng/ml for CT26), separately, for 48 hours. Live/dead cells were stained with a Calcein/Propidium Iodide (PI) cell viability/cytotoxicity assay kit (Beyotime, #C2015) for 30 min at 37°C. Cells were observed under a fluorescence microscope (Leica). The number of fluorescent cells was counted with ImageJ (https://imagej.nih.gov/ij/). Difference in the numbers of live/dead cells between control and treated cells were analyzed with unpaired Student’s *t*-test based on six replicates.

## Notes

https://www.ncbi.nlm.nih.gov/geo/query/acc.cgi?acc=GSE298039

https://www.ncbi.nlm.nih.gov/geo/query/acc.cgi?acc=GSE295152

https://www.ncbi.nlm.nih.gov/geo/query/acc.cgi?acc=GSE295153

https://www.ncbi.nlm.nih.gov/geo/query/acc.cgi?acc=GSE295154

https://www.ncbi.nlm.nih.gov/geo/query/acc.cgi?acc=GSE295155

https://www.ncbi.nlm.nih.gov/geo/query/acc.cgi?acc=GSE295156

https://www.ncbi.nlm.nih.gov/geo/query/acc.cgi?acc=GSE295157

https://www.ncbi.nlm.nih.gov/geo/query/acc.cgi?acc=GSE295158

https://www.ncbi.nlm.nih.gov/geo/query/acc.cgi?acc=GSE295159

### Competing Interest Statement

The authors have declared no competing interest.

### Funding

- Shenzhen Science and Technology Program: JCYJ20210324120205015

## References

Agudo J, Miao Y. Stemness in solid malignancies: coping with immune attack. Nat Rev Cancer. 2025 Jan;25(1):27–40.

Ahmad F, Patrick S, Sheikh T, Sharma V, Pathak P, Malgulwar PB, Kumar A, Joshi SD, Sarkar C, Sen E. Telomerase reverse transcriptase (TERT) - enhancer of zeste homolog 2 (EZH2) network regulates lipid metabolism and DNA damage responses in glioblastoma. J Neurochem. 2017 Dec;143(6):671–683.

Ascic E, Åkerström F, Sreekumar Nair M, Rosa A, Kurochkin I, Zimmermannova O, Catena X, Rotankova N, Veser C, Rudnik M, Ballocci T, Schärer T, Huang X, de Rosa Torres M, Renaud E, Velasco Santiago M, Met Ö, Askmyr D, Lindstedt M, Greiff L, Ligeon LA, Agarkova I, Svane IM, Pires CF, Rosa FF, Pereira CF. In vivo dendritic cell reprogramming for cancer immunotherapy. Science. 2024 Oct 18;386(6719):eadn9083.

Balinth S, Fisher ML, Hwangbo Y, Wu C, Ballon C, Sun X, Mills AA. EZH2 regulates a SETDB1/ΔNp63α axis via RUNX3 to drive a cancer stem cell phenotype in squamous cell carcinoma. Oncogene. 2022 Aug;41(35):4130–4144.

Bryant RJ, Cross NA, Eaton CL, Hamdy FC, Cunliffe VT. EZH2 promotes proliferation and invasiveness of prostate cancer cells. Prostate. 2007 Apr 1;67(5):547–56.

Cao Y. Tumorigenesis as a process of gradual loss of original cell identity and gain of properties of neural precursor/progenitor cells. Cell Biosci. 2017 Nov 7;7:61.

Cao Y. Neural is Fundamental: Neural Stemness as the Ground State of Cell Tumorigenicity and Differentiation Potential. Stem Cell Rev Rep. 2022 Jan;18(1):37–55.

Cao Y. Neural induction drives body axis formation during embryogenesis, but a neural induction-like process drives tumorigenesis in postnatal animals. Front Cell Dev Biol. 2023 May 9;11:1092667.

Cao Y. Lack of basic rationale in epithelial-mesenchymal transition and its related concepts. Cell Biosci. 2024 Aug 20;14(1):104.

Champiat S, Ferrara R, Massard C, Besse B, Marabelle A, Soria JC, Ferté C. Hyperprogressive disease: recognizing a novel pattern to improve patient management. Nat Rev Clin Oncol. 2018 Dec;15(12):748–762.

Chen L, Zhang M, Fang L, Yang X, Cao N, Xu L, Shi L, Cao Y. Coordinated regulation of the ribosome and proteasome by PRMT1 in the maintenance of neural stemness in cancer cells and neural stem cells. J Biol Chem. 2021 Nov;297(5):101275.

Crea F, Paolicchi E, Marquez VE, Danesi R. Polycomb genes and cancer: time for clinical application? Crit Rev Oncol Hematol. 2012 Aug;83(2):184–93.

de Miguel M, Calvo E. Clinical Challenges of Immune Checkpoint Inhibitors. Cancer Cell. 2020 Sep 14;38(3):326–333.

Deng J, Wang ES, Jenkins RW, Li S, Dries R, Yates K, Chhabra S, Huang W, Liu H, Aref AR, Ivanova E, Paweletz CP, Bowden M, Zhou CW, Herter-Sprie GS, Sorrentino JA, Bisi JE, Lizotte PH, Merlino AA, Quinn MM, Bufe LE, Yang A, Zhang Y, Zhang H, Gao P, Chen T, Cavanaugh ME, Rode AJ, Haines E, Roberts PJ, Strum JC, Richards WG, Lorch JH, Parangi S, Gunda V, Boland GM, Bueno R, Palakurthi S, Freeman GJ, Ritz J, Haining WN, Sharpless NE, Arthanari H, Shapiro GI, Barbie DA, Gray NS, Wong KK. CDK4/6 Inhibition Augments Antitumor Immunity by Enhancing T-cell Activation. Cancer Discov. 2018 Feb;8(2):216–233.

De Sá Fernandes C, Novoszel P, Gastaldi T, Krauß D, Lang M, Rica R, Kutschat AP, Holcmann M, Ellmeier W, Seruggia D, Strobl H, Sibilia M. The histone deacetylase HDAC1 controls dendritic cell development and anti-tumor immunity. Cell Rep. 2024 Jun 25;43(6):114308.

de Thé H. Differentiation therapy revisited. Nat Rev Cancer. 2018 Feb;18(2):117–127.

Dhanasekaran R, Deutzmann A, Mahauad-Fernandez WD, Hansen AS, Gouw AM, Felsher DW. The MYC oncogene - the grand orchestrator of cancer growth and immune evasion. Nat Rev Clin Oncol. 2022 Jan;19(1):23–36.

Draghi A, Chamberlain CA, Furness A, Donia M. Acquired resistance to cancer immunotherapy. Semin Immunopathol. 2019 Jan;41(1):31–40.

Drukker M, Katchman H, Katz G, Even-Tov Friedman S, Shezen E, Hornstein E, Mandelboim O, Reisner Y, Benvenisty N. Human embryonic stem cells and their differentiated derivatives are less susceptible to immune rejection than adult cells. Stem Cells. 2006 Feb;24(2):221–9.

Fändrich F, Dresske B, Bader M, Schulze M. Embryonic stem cells share immune-privileged features relevant for tolerance induction. J Mol Med (Berl). 2002 Jun;80(6):343–50.

Fatma H, Maurya SK, Siddique HR. Epigenetic modifications of c-MYC: Role in cancer cell reprogramming, progression and chemoresistance. Semin Cancer Biol. 2022 Aug;83:166–176.

Galassi C, Musella M, Manduca N, Maccafeo E, Sistigu A. The Immune Privilege of Cancer Stem Cells: A Key to Understanding Tumor Immune Escape and Therapy Failure. Cells. 2021 Sep 8;10(9):2361.

Garlanda C, Mantovani A. Interleukin-1 in tumor progression, therapy, and prevention. Cancer Cell. 2021 Aug 9;39(8):1023–1027.

Gide TN, Wilmott JS, Scolyer RA, Long GV. Primary and Acquired Resistance to Immune Checkpoint Inhibitors in Metastatic Melanoma. Clin Cancer Res. 2018 Mar 15;24(6):1260–1270.

Gong JR, Lee CK, Kim HM, Kim J, Jeon J, Park S, Cho KH. Control of Cellular Differentiation Trajectories for Cancer Reversion. Adv Sci (Weinh). 2025 Jan;12(3):e2402132.

Griffin GK, Wu J, Iracheta-Vellve A, Patti JC, Hsu J, Davis T, Dele-Oni D, Du PP, Halawi AG, Ishizuka JJ, Kim SY, Klaeger S, Knudsen NH, Miller BC, Nguyen TH, Olander KE, Papanastasiou M, Rachimi S, Robitschek EJ, Schneider EM, Yeary MD, Zimmer MD, Jaffe JD, Carr SA, Doench JG, Haining WN, Yates KB, Manguso RT, Bernstein BE. Epigenetic silencing by SETDB1 suppresses tumour intrinsic immunogenicity. Nature. 2021 Jul;595(7866):309–314.

He L, Azizad D, Bhat K, Ioannidis A, Hoffmann CJ, Arambula E, Eghbali M, Bhaduri A, Kornblum HI, Pajonk F. Radiation-induced cellular plasticity primes glioblastoma for forskolin-mediated differentiation. Proc Natl Acad Sci U S A. 2025 Mar 4;122(9):e2415557122.

Hori J, Ng TF, Shatos M, Klassen H, Streilein JW, Young MJ. Neural progenitor cells lack immunogenicity and resist destruction as allografts. Stem Cells. 2003;21(4):405–16.

Huang L, Malu S, McKenzie JA, Andrews MC, Talukder AH, Tieu T, Karpinets T, Haymaker C, Forget MA, Williams LJ, Wang Z, Mbofung RM, Wang ZQ, Davis RE, Lo RS, Wargo JA, Davies MA, Bernatchez C, Heffernan T, Amaria RN, Korkut A, Peng W, Roszik J, Lizée G, Woodman SE, Hwu P. The RNA-binding Protein MEX3B Mediates Resistance to Cancer Immunotherapy by Downregulating HLA-A Expression. Clin Cancer Res. 2018 Jul 15;24(14):3366–3376.

Ishizuka JJ, Manguso RT, Cheruiyot CK, Bi K, Panda A, Iracheta-Vellve A, Miller BC, Du PP, Yates KB, Dubrot J, Buchumenski I, Comstock DE, Brown FD, Ayer A, Kohnle IC, Pope HW, Zimmer MD, Sen DR, Lane-Reticker SK, Robitschek EJ, Griffin GK, Collins NB, Long AH, Doench JG, Kozono D, Levanon EY, Haining WN. Loss of ADAR1 in tumours overcomes resistance to immune checkpoint blockade. Nature. 2019 Jan;565(7737):43–48.

Itakura G, Ozaki M, Nagoshi N, Kawabata S, Nishiyama Y, Sugai K, Iida T, Kashiwagi R, Ookubo T, Yastake K, Matsubayashi K, Kohyama J, Iwanami A, Matsumoto M, Nakamura M, Okano H. Low immunogenicity of mouse induced pluripotent stem cell-derived neural stem/progenitor cells. Sci Rep. 2017 Oct 11;7(1):12996.

Joyce JA, Fearon DT. T cell exclusion, immune privilege, and the tumor microenvironment. Science. 2015 Apr 3;348(6230):74–80.

Jurcak NR, Rucki AA, Muth S, Thompson E, Sharma R, Ding D, Zhu Q, Eshleman JR, Anders RA, Jaffee EM, Fujiwara K, Zheng L. Axon Guidance Molecules Promote Perineural Invasion and Metastasis of Orthotopic Pancreatic Tumors in Mice. Gastroenterology. 2019 Sep;157(3):838–850.e6.

Kalbasi A, Ribas A. Tumour-intrinsic resistance to immune checkpoint blockade. Nat Rev Immunol. 2020 Jan;20(1):25–39.

Kamada T, Togashi Y, Tay C, Ha D, Sasaki A, Nakamura Y, Sato E, Fukuoka S, Tada Y, Tanaka A, Morikawa H, Kawazoe A, Kinoshita T, Shitara K, Sakaguchi S, Nishikawa H. PD-1+ regulatory T cells amplified by PD-1 blockade promote hyperprogression of cancer. Proc Natl Acad Sci U S A. 2019 May 14;116(20):9999–10008.

Karasarides M, Cogdill AP, Robbins PB, Bowden M, Burton EM, Butterfield LH, Cesano A, Hammer C, Haymaker CL, Horak CE, McGee HM, Monette A, Rudqvist NP, Spencer CN, Sweis RF, Vincent BG, Wennerberg E, Yuan J, Zappasodi R, Lucey VMH, Wells DK, LaVallee T. Hallmarks of Resistance to Immune-Checkpoint Inhibitors. Cancer Immunol Res. 2022 Apr 1;10(4):372–383.

Kearney CJ, Vervoort SJ, Hogg SJ, Ramsbottom KM, Freeman AJ, Lalaoui N, Pijpers L, Michie J, Brown KK, Knight DA, Sutton V, Beavis PA, Voskoboinik I, Darcy PK, Silke J, Trapani JA, Johnstone RW, Oliaro J. Tumor immune evasion arises through loss of TNF sensitivity. Sci Immunol. 2018 May 18;3(23):eaar3451.

Kerk SA, Papagiannakopoulos T, Shah YM, Lyssiotis CA. Metabolic networks in mutant KRAS-driven tumours: tissue specificities and the microenvironment. Nat Rev Cancer. 2021 Aug;21(8):510–525.

Kim HJ, Cantor H, Cosmopoulos K. Overcoming Immune Checkpoint Blockade Resistance via EZH2 Inhibition. Trends Immunol. 2020 Oct;41(10):948–963.

Krah NM, Narayanan SM, Yugawa DE, Straley JA, Wright CVE, MacDonald RJ, Murtaugh LC. Prevention and Reversion of Pancreatic Tumorigenesis through a Differentiation-Based Mechanism. Dev Cell. 2019 Sep 23;50(6):744–754.e4.

Lasse-Opsahl EL, Barravecchia I, McLintock E, Lee JM, Ferris SF, Espinoza CE, Hinshaw R, Cavanaugh S, Robotti M, Rober L, Brown K, Abdelmalak KY, Galban CJ, Frankel TL, Zhang Y, Pasca di Magliano M, Galban S. KRASG12D drives immunosuppression in lung adenocarcinoma through paracrine signaling. JCI Insight. 2025 Jan 9;10(1):e182228.

Lee SH, Jeyapalan JN, Appleby V, Mohamed Noor DA, Sottile V, Scotting PJ. Dynamic methylation and expression of Oct4 in early neural stem cells. J Anat. 2010 Sep;217(3):203–13.

Lei A, Chen L, Zhang M, Yang X, Xu L, Cao N, Zhang Z, Cao Y. EZH2 Regulates Protein Stability via Recruiting USP7 to Mediate Neuronal Gene Expression in Cancer Cells. Front Genet. 2019 May 3;10:422.

Li J, Stanger BZ. How Tumor Cell Dedifferentiation Drives Immune Evasion and Resistance to Immunotherapy. Cancer Res. 2020 Oct 1;80(19):4037–4041.

Linde MH, Fan AC, Köhnke T, Trotman-Grant AC, Gurev SF, Phan P, Zhao F, Haddock NL, Nuno KA, Gars EJ, Stafford M, Marshall PL, Dove CG, Linde IL, Landberg N, Miller LP, Majzner RG, Zhang TY, Majeti R. Reprogramming Cancer into Antigen-Presenting Cells as a Novel Immunotherapy. Cancer Discov. 2023 May 4;13(5):1164–1185.

Llombart V, Mansour MR. Therapeutic targeting of “undruggable” MYC. EBioMedicine. 2022 Jan;75:103756.

Lupo F, Pezzini F, Pasini D, Fiorini E, Adamo A, Veghini L, Bevere M, Frusteri C, Delfino P, D’agosto S, Andreani S, Piro G, Malinova A, Wang T, De Sanctis F, Lawlor RT, Hwang CI, Carbone C, Amelio I, Bailey P, Bronte V, Tuveson D, Scarpa A, Ugel S, Corbo V. Axon guidance cue SEMA3A promotes the aggressive phenotype of basal-like PDAC. Gut. 2024 Jul 11;73(8):1321–1335.

Magliocca JF, Held IK, Odorico JS. Undifferentiated murine embryonic stem cells cannot induce portal tolerance but may possess immune privilege secondary to reduced major histocompatibility complex antigen expression. Stem Cells Dev. 2006 Oct;15(5):707–17.

Mahadevan KK, McAndrews KM, LeBleu VS, Yang S, Lyu H, Li B, Sockwell AM, Kirtley ML, Morse SJ, Moreno Diaz BA, Kim MP, Feng N, Lopez AM, Guerrero PA, Paradiso F, Sugimoto H, Arian KA, Ying H, Barekatain Y, Sthanam LK, Kelly PJ, Maitra A, Heffernan TP, Kalluri R. KRASG12D inhibition reprograms the microenvironment of early and advanced pancreatic cancer to promote FAS-mediated killing by CD8+ T cells. Cancer Cell. 2023 Sep 11;41(9):1606–1620.e8.

Marcucci F, Rumio C. The tumor-promoting effects of the adaptive immune system: a cause of hyperprogressive disease in cancer? Cell Mol Life Sci. 2021 Feb;78(3):853–865.

Miao Y, Yang H, Levorse J, Yuan S, Polak L, Sribour M, Singh B, Rosenblum MD, Fuchs E. Adaptive Immune Resistance Emerges from Tumor-Initiating Stem Cells. Cell. 2019 May 16;177(5):1172–1186.e14.

Miller DM, Thomas SD, Islam A, Muench D, Sedoris K. c-Myc and cancer metabolism. Clin Cancer Res. 2012 Oct 15;18(20):5546–53.

Nylund P, Atienza Párraga A, Haglöf J, De Bruyne E, Menu E, Garrido-Zabala B, Ma A, Jin J, Öberg F, Vanderkerken K, Kalushkova A, Jernberg-Wiklund H. A distinct metabolic response characterizes sensitivity to EZH2 inhibition in multiple myeloma. Cell Death Dis. 2021 Feb 12;12(2):167.

O’Donnell JS, Teng MWL, Smyth MJ. Cancer immunoediting and resistance to T cell-based immunotherapy. Nat Rev Clin Oncol. 2019 Mar;16(3):151–167.

Ougolkov AV, Bilim VN, Billadeau DD. Regulation of pancreatic tumor cell proliferation and chemoresistance by the histone methyltransferase enhancer of zeste homologue 2. Clin Cancer Res. 2008 Nov 1;14(21):6790–6.

Ozaki M, Iwanami A, Nagoshi N, Kohyama J, Itakura G, Iwai H, Nishimura S, Nishiyama Y, Kawabata S, Sugai K, Iida T, Matsubayashi K, Isoda M, Kashiwagi R, Toyama Y, Matsumoto M, Okano H, Nakamura M. Evaluation of the immunogenicity of human iPS cell-derived neural stem/progenitor cells in vitro. Stem Cell Res. 2017 Mar;19:128–138.

Pierce GB, Wallace C. Differentiation of malignant to benign cells. Cancer Res. 1971 Feb;31(2):127–34.

Rathjen J, Haines BP, Hudson KM, Nesci A, Dunn S, Rathjen PD. Directed differentiation of pluripotent cells to neural lineages: homogeneous formation and differentiation of a neurectoderm population. Development. 2002 Jun;129(11):2649–61.

Ruiz de Galarreta M, Bresnahan E, Molina-Sánchez P, Lindblad KE, Maier B, Sia D, Puigvehi M, Miguela V, Casanova-Acebes M, Dhainaut M, Villacorta-Martin C, Singhi AD, Moghe A, von Felden J, Tal Grinspan L, Wang S, Kamphorst AO, Monga SP, Brown BD, Villanueva A, Llovet JM, Merad M, Lujambio A. β-Catenin Activation Promotes Immune Escape and Resistance to Anti-PD-1 Therapy in Hepatocellular Carcinoma. Cancer Discov. 2019 Aug;9(8):1124–1141.

Schade AE, Perurena N, Yang Y, Rodriguez CL, Krishnan A, Gardner A, Loi P, Xu Y, Nguyen VTM, Mastellone GM, Pilla NF, Watanabe M, Ota K, Davis RA, Mattioli K, Xiang D, Zoeller JJ, Lin JR, Morganti S, Garrido-Castro AC, Tolaney SM, Li Z, Barbie DA, Sorger PK, Helin K, Santagata S, Knott SRV, Cichowski K. AKT and EZH2 inhibitors kill TNBCs by hijacking mechanisms of involution. Nature. 2024 Nov;635(8039):755–763.

Schoenfeld AJ, Hellmann MD. Acquired Resistance to Immune Checkpoint Inhibitors. Cancer Cell. 2020 Apr 13;37(4):443–455.

Shah NN, Fry TJ. Mechanisms of resistance to CAR T cell therapy. Nat Rev Clin Oncol. 2019;16(6):372–385.

Sharma P, Hu-Lieskovan S, Wargo JA, Ribas A. Primary, Adaptive, and Acquired Resistance to Cancer Immunotherapy. Cell. 2017 Feb 9;168(4):707–723.

Solter D. From teratocarcinomas to embryonic stem cells and beyond: a history of embryonic stem cell research. Nat Rev Genet. 2006 Apr;7(4):319–27.

Spranger S, Bao R, Gajewski TF. Melanoma-intrinsic β-catenin signalling prevents anti-tumour immunity. Nature. 2015 Jul 9;523(7559):231-5.

van Loo G, Bertrand MJM. Death by TNF: a road to inflammation. Nat Rev Immunol. 2023 May;23(5):289–303.

Vinay DS, Ryan EP, Pawelec G, Talib WH, Stagg J, Elkord E, Lichtor T, Decker WK, Whelan RL, Kumara HMCS, Signori E, Honoki K, Georgakilas AG, Amin A, Helferich WG, Boosani CS, Guha G, Ciriolo MR, Chen S, Mohammed SI, Azmi AS, Keith WN, Bilsland A, Bhakta D, Halicka D, Fujii H, Aquilano K, Ashraf SS, Nowsheen S, Yang X, Choi BK, Kwon BS. Immune evasion in cancer: Mechanistic basis and therapeutic strategies. Semin Cancer Biol. 2015 Dec;35 Suppl:S185–S198.

Vitale I, Shema E, Loi S, Galluzzi L. Intratumoral heterogeneity in cancer progression and response to immunotherapy. Nat Med. 2021 Feb;27(2):212–224.

Watterson A, Coelho MA. Cancer immune evasion through KRAS and PD-L1 and potential therapeutic interventions. Cell Commun Signal. 2023 Mar 2;21(1):45.

Xu L, Zhang M, Shi L, Yang X, Chen L, Cao N, Lei A, Cao Y. Neural stemness contributes to cell tumorigenicity. Cell Biosci. 2021 Jan 19;11(1):21.

Yang J, Antin P, Berx G, Blanpain C, Brabletz T, Bronner M, Campbell K, Cano A, Casanova J, Christofori G, Dedhar S, Derynck R, Ford HL, Fuxe J, García de Herreros A, Goodall GJ, Hadjantonakis AK, Huang RYJ, Kalcheim C, Kalluri R, Kang Y, Khew-Goodall Y, Levine H, Liu J, Longmore GD, Mani SA, Massagué J, Mayor R, McClay D, Mostov KE, Newgreen DF, Nieto MA, Puisieux A, Runyan R, Savagner P, Stanger B, Stemmler MP, Takahashi Y, Takeichi M, Theveneau E, Thiery JP, Thompson EW, Weinberg RA, Williams ED, Xing J, Zhou BP, Sheng G; EMT International Association (TEMTIA). Guidelines and definitions for research on epithelial-mesenchymal transition. Nat Rev Mol Cell Biol. 2020 Jun;21(6):341–352.

Yang X, Cao N, Chen L, Liu L, Zhang M, Cao Y. Suppression of Cell Tumorigenicity by Non-neural Pro-differentiation Factors via Inhibition of Neural Property in Tumorigenic Cells. Front Cell Dev Biol. 2021 Sep 14;9:714383.

Zhang M, Liu Y, Shi L, Fang L, Xu L, Cao Y. Neural stemness unifies cell tumorigenicity and pluripotent differentiation potential. J Biol Chem. 2022 Jul;298(7):102106.

Zhang Z, Lei A, Xu L, Chen L, Chen Y, Zhang X, Gao Y, Yang X, Zhang M, Cao Y. Similarity in gene-regulatory networks suggests that cancer cells share characteristics of embryonic neural cells. J Biol Chem. 2017 Aug 4;292(31):12842–12859.

Zhong H, Lu W, Tang Y, Wiel C, Wei Y, Cao J, Riedlinger G, Papagiannakopoulos T, Guo JY, Bergo MO, Kang Y, Ganesan S, Sabaawy HE, Pine SR. SOX9 drives KRAS-induced lung adenocarcinoma progression and suppresses anti-tumor immunity. Oncogene. 2023 Jun;42(27):2183–2194.

Zhou L, Mudianto T, Ma X, Riley R, Uppaluri R. Targeting EZH2 Enhances Antigen Presentation, Antitumor Immunity, and Circumvents Anti-PD-1 Resistance in Head and Neck Cancer. Clin Cancer Res. 2020 Jan 1;26(1):290–300.

Zimmerli D, Brambillasca CS, Talens F, Bhin J, Linstra R, Romanens L, Bhattacharya A, Joosten SEP, Da Silva AM, Padrao N, Wellenstein MD, Kersten K, de Boo M, Roorda M, Henneman L, de Bruijn R, Annunziato S, van der Burg E, Drenth AP, Lutz C, Endres T, van de Ven M, Eilers M, Wessels L, de Visser KE, Zwart W, Fehrmann RSN, van Vugt MATM, Jonkers J. MYC promotes immune-suppression in triple-negative breast cancer via inhibition of interferon signaling. Nat Commun. 2022 Nov 2;13(1):6579.

Zimmermannova O, Ferreira AG, Ascic E, Velasco Santiago M, Kurochkin I, Hansen M, Met Ö, Caiado I, Shapiro IE, Michaux J, Humbert M, Soto-Cabrera D, Benonisson H, Silvério-Alves R, Gomez-Jimenez D, Bernardo C, Bauden M, Andersson R, Höglund M, Miharada K, Nakamura Y, Hugues S, Greiff L, Lindstedt M, Rosa FF, Pires CF, Bassani-Sternberg M, Svane IM, Pereira CF. Restoring tumor immunogenicity with dendritic cell reprogramming. Sci Immunol. 2023 Jul 14;8(85):eadd4817.

Zingg D, Debbache J, Schaefer SM, Tuncer E, Frommel SC, Cheng P, Arenas-Ramirez N, Haeusel J, Zhang Y, Bonalli M, McCabe MT, Creasy CL, Levesque MP, Boyman O, Santoro R, Shakhova O, Dummer R, Sommer L. The epigenetic modifier EZH2 controls melanoma growth and metastasis through silencing of distinct tumour suppressors. Nat Commun. 2015 Jan 22;6:6051.

