## Supplemental data for "Differentiation status determines tumorigenicity and immunogenicity of cancer cells"

### Supplemental Figures 1-9 and figure legends

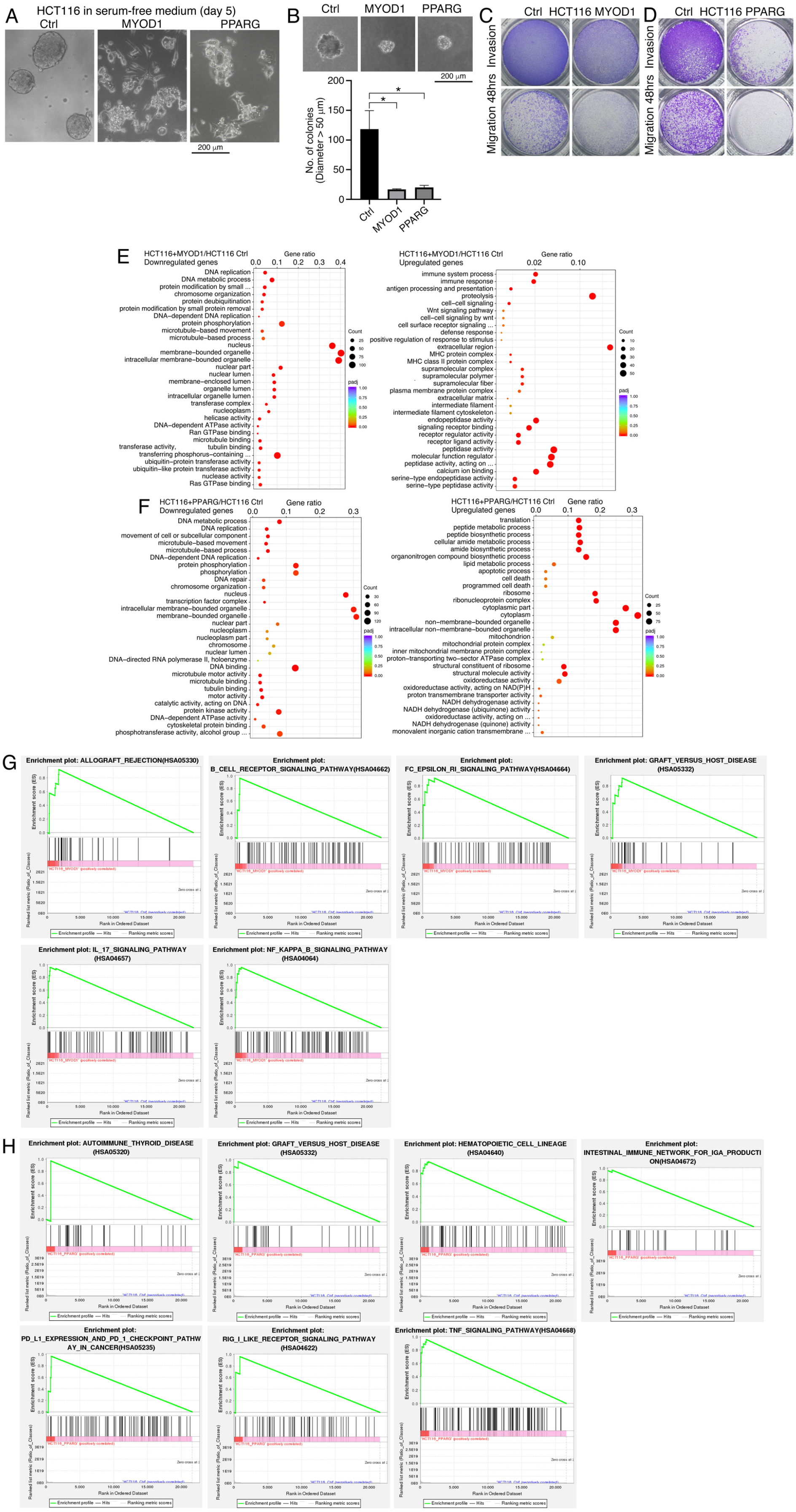

**Fig. S1. Changes in cellular properties and transcriptomes of HCT116 cells after induced differentiation.** (A-C) Cellular property change in control (Ctrl) and cells with forced expression of MYOD1 or PPARG, as shown by neurosphere formation in NSC-specific serum-free medium (A), colony formation in soft agar (B), and invasion and migration in transwell assays (C, D). In (B), significance of difference in colony formation was calculated based on experiments in triplicate using unpaired Student's *t*-test. Data are shown as mean±SEM. *p < 0.05. Colonies with a diameter > 50 µm were counted. (E-H) Change in transcriptomes in cells after induced differentiation, as compared with the transcriptome of control cells. GO (E, F) analyses show enrichment of downregulated and upregulated genes, and GSEA (G, H) show enrichment of immune related gene sets in cells with forced expression of MYOD1 (E, G) or PPARG (F, H).

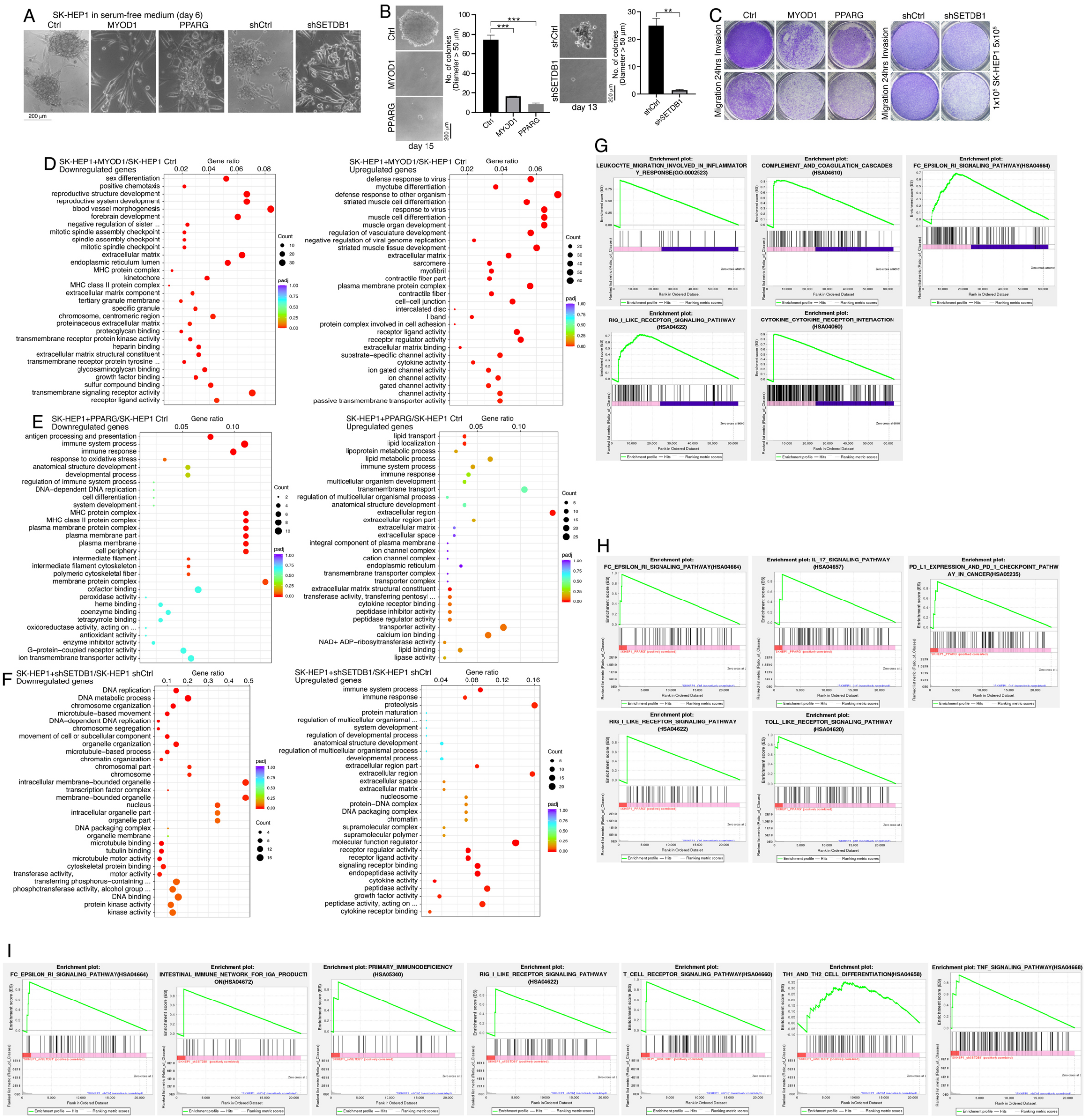

**Fig. S2. Changes in cellular properties and transcriptomes of SK-HEP1 cells after induced differentiation.** (A-C) Cellular property change in control (Ctrl, shCtrl) and cells with forced expression of MYOD1 or PPARG, or knockdown of SETDB1, as shown by neurosphere formation in NSC-specific serum-free medium (A), colony formation in soft agar (B), and invasion and migration in transwell assays (C, D). In (B), significance of difference in colony formation was calculated based on experiments in triplicate using unpaired Student's *t*-test. Data are shown as mean±SEM. *p < 0.05. Colonies with a diameter > 50 μm were counted. (D-I) Comparison of transcriptomes of cells with forced expression of MYOD1 or PPARG, or knockdown of SETDB1 with the transcriptome of control cells. GO (D-F) analyses show enrichment of downregulated and upregulated genes, and GSEA (G-I) show enrichment of immune related gene sets in cells with forced expression of MYOD1 (D, G) or PPARG (E, H), or knockdown of SETDB1 (F, I).

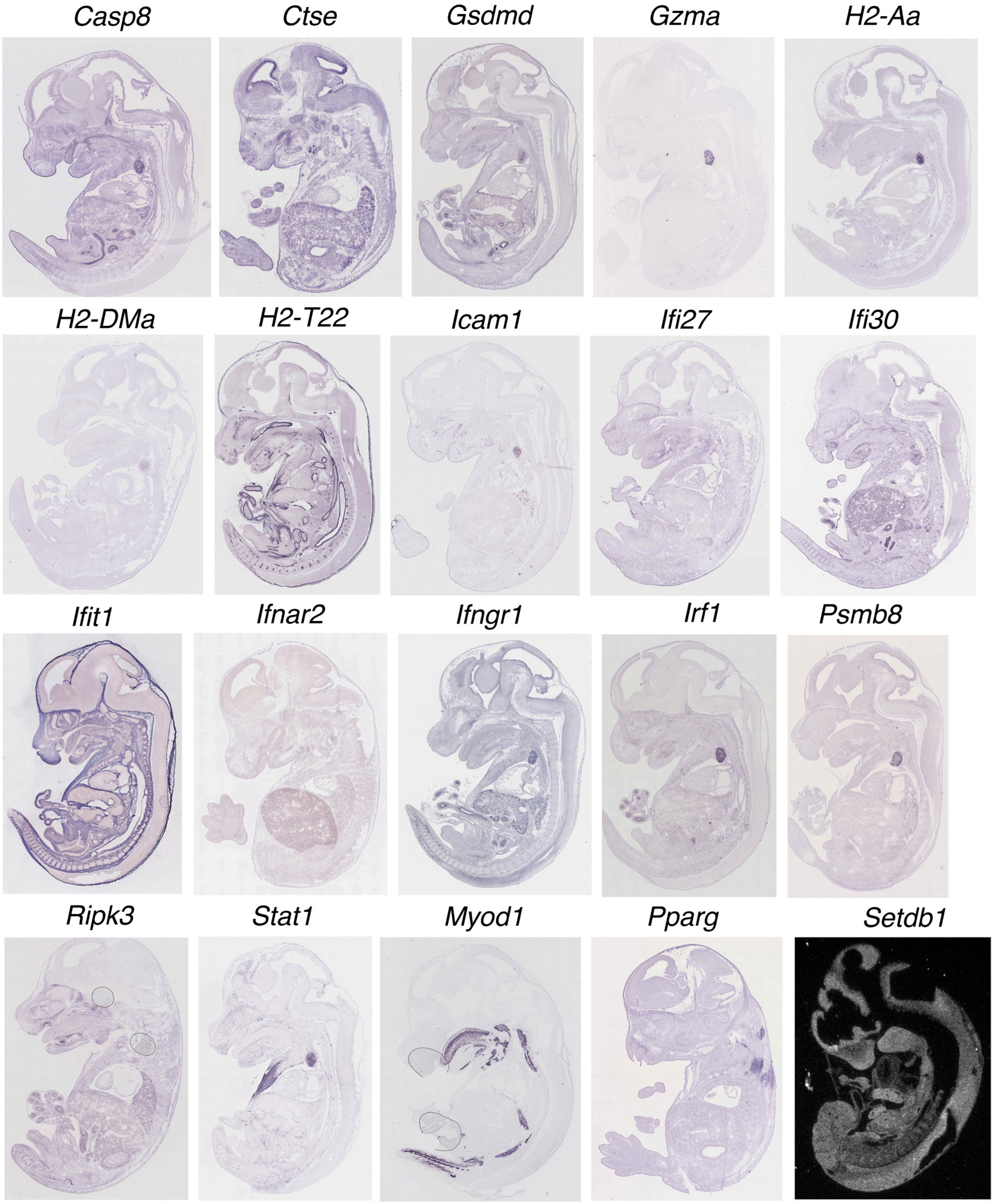


**Fig. S3. Comparison of spatial expression patterns of immune related genes involved in Ifn-γ response, antigen processing and presentation, Tnf-α mediated cell death, lineage differentiation, and neural stemness genes**. Major tissues where the genes are expressed are as follows (for details, refer to https://www.informatics.jax.org/). *Casp8*: thymus primordium; *Ctse*: liver, lung, submandibular gland primordium, vibrissa, dorsal root ganglion, etc.; *Gsdmd*: midgut, hindgut, thymus primordium; *Gzma*: thymus primordium; *H2-Aa*: thymus primordium; *H2-DMa*: thymus primordium; *H2-T22*: vertebral axis musculature, limb, stomach, smooth muscle tissue, gut, esophagus, etc.; *Icam1*: lung, thymus primordium; *Ifi27*: testis, adrenal gland; *Ifi30*: liver, orbito-sphenoid, stomach, gut, esophagus, thymus primordium, etc.; *Ifit1*: liver, cardiovascular system, skin, limb, integumental system, etc.; *Ifnar2*: liver; *Ifngr1*: chondrocranium, liver, Meckel's cartilage, jaw molar, thymus primordium, etc.; *Irf1*: testis, thymus primordium, midgut; *Psmb8*: thymus primordium; *Ripk3*: jaw molar, thymus primordium, turbinate bone primordium; *Stat1*: axial musculature, pectoral girdle and thoracic body wall muscle, thymus primordium, etc.; *Myod1*: vertebral axis musculature, diaphragm, mesenchymes, tongue muscle; *Pparg*: axial musculature, bladder fundus, urethra; *Setdb1*: nervous system. Expression patterns of all genes except *Setdb1* were generated with RNA in situ hybridization on cryosections of E14.5 mouse embryos. Expression pattern of *Setdb1* was obtained with radio-labeled RNA in situ hybridization on sections of E11.5 embryo and visualized with autoradiography. All images were reproduced from https://www.informatics.jax.org/ under the CC BY 4.0 license.

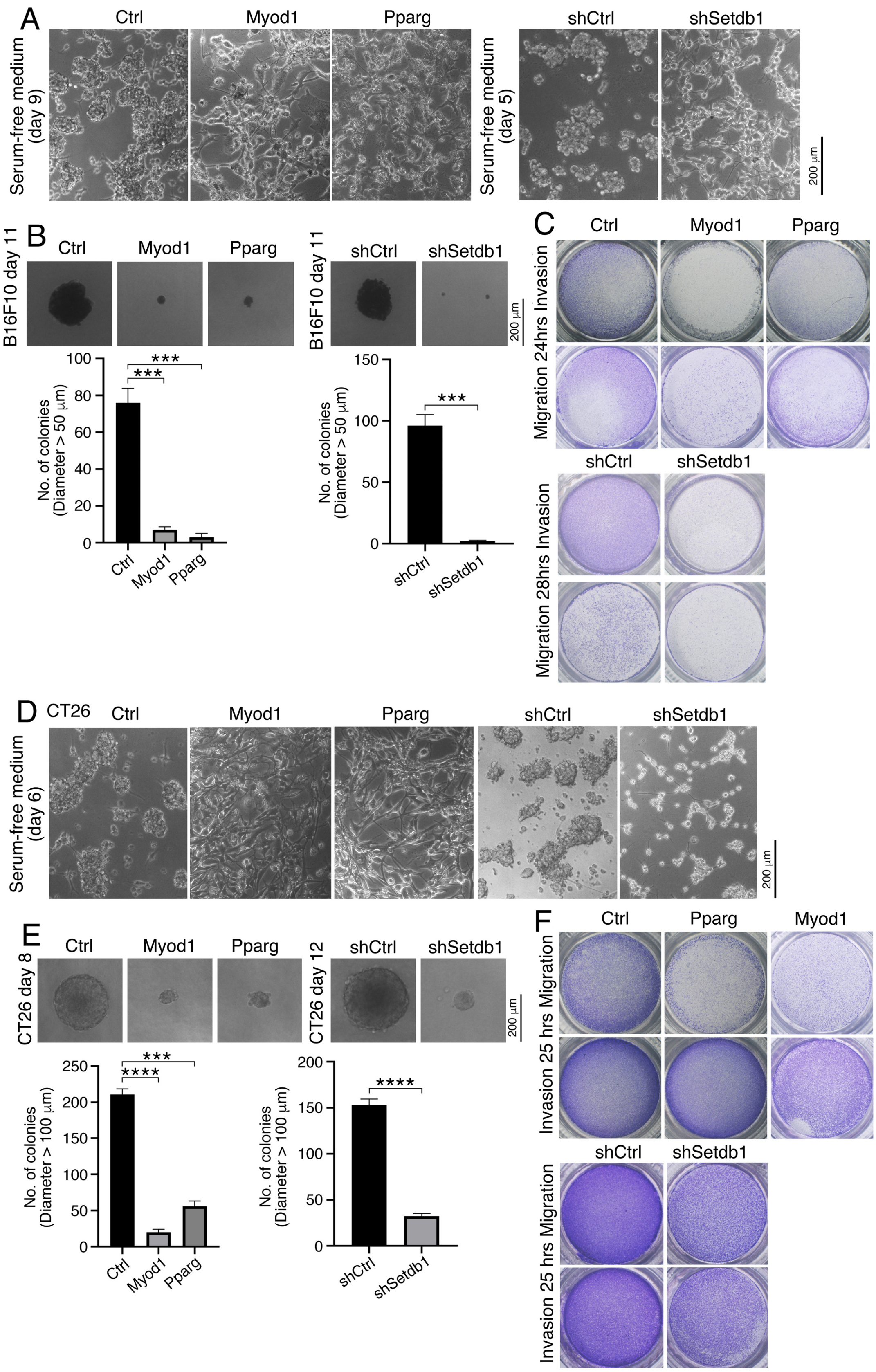


**Fig. S4. Cellular property alteration after differentiation of B16F10 or CT26 cells.** Characterization of cellular properties were performed by culturing cells in NSC-specific serum-free medium (A, D), colony formation in soft agar (B, E), and invasion and migration assays (C, F) in control (Ctrl, shCtrl) and differentiated B16F10 (A-C) or CT26 (D-F) cells in response to forced expression of Myod1, Pparg, or knockdown of Setdb1 (shSetdb1). In (B) and (E), significance of difference in colony formation was calculated based on experiments in triplicate using unpaired Student's *t*-test. Data are shown as mean±SEM. ***$p < 0.001$, ****$p < 0.0001$. In (B), colonies with a diameter > 50 μm were counted; in (E), colonies with a diameter > 100 μm were counted.

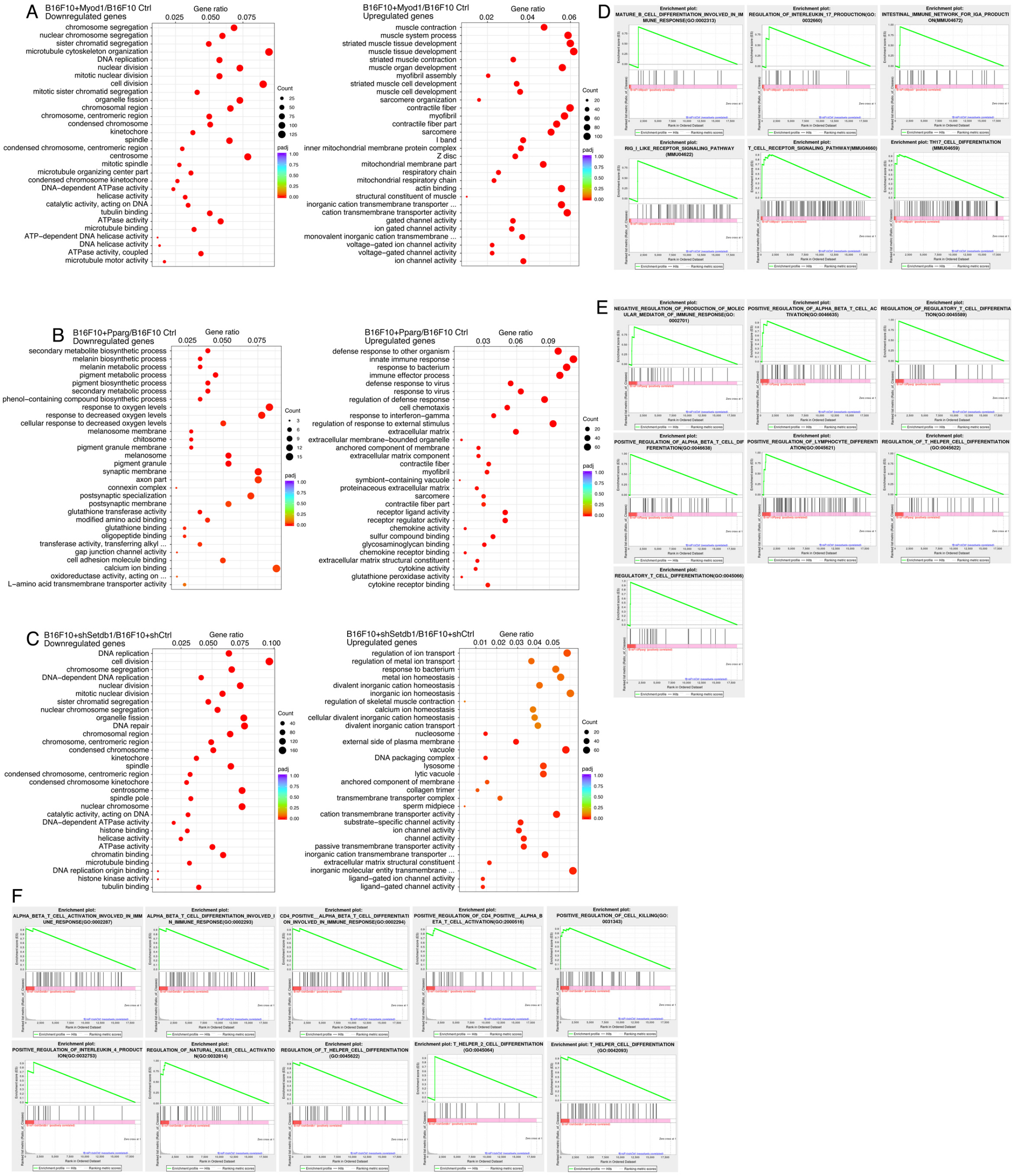


**Fig. S5. Transcriptome profiling of control B16F10 cells and cells after induced differentiation.** (A-C) GO analysis on downregulated and upregulated genes in differentiated cells with forced expression of Myod1 (A), Pparg (B), or Setdb1 knockdown (C). (D-F) Immune related gene sets enriched by GSEA in differentiated cells in response to forced expression of Myod1 (D), Pparg (E), or knockdown of Setdb1 (F).

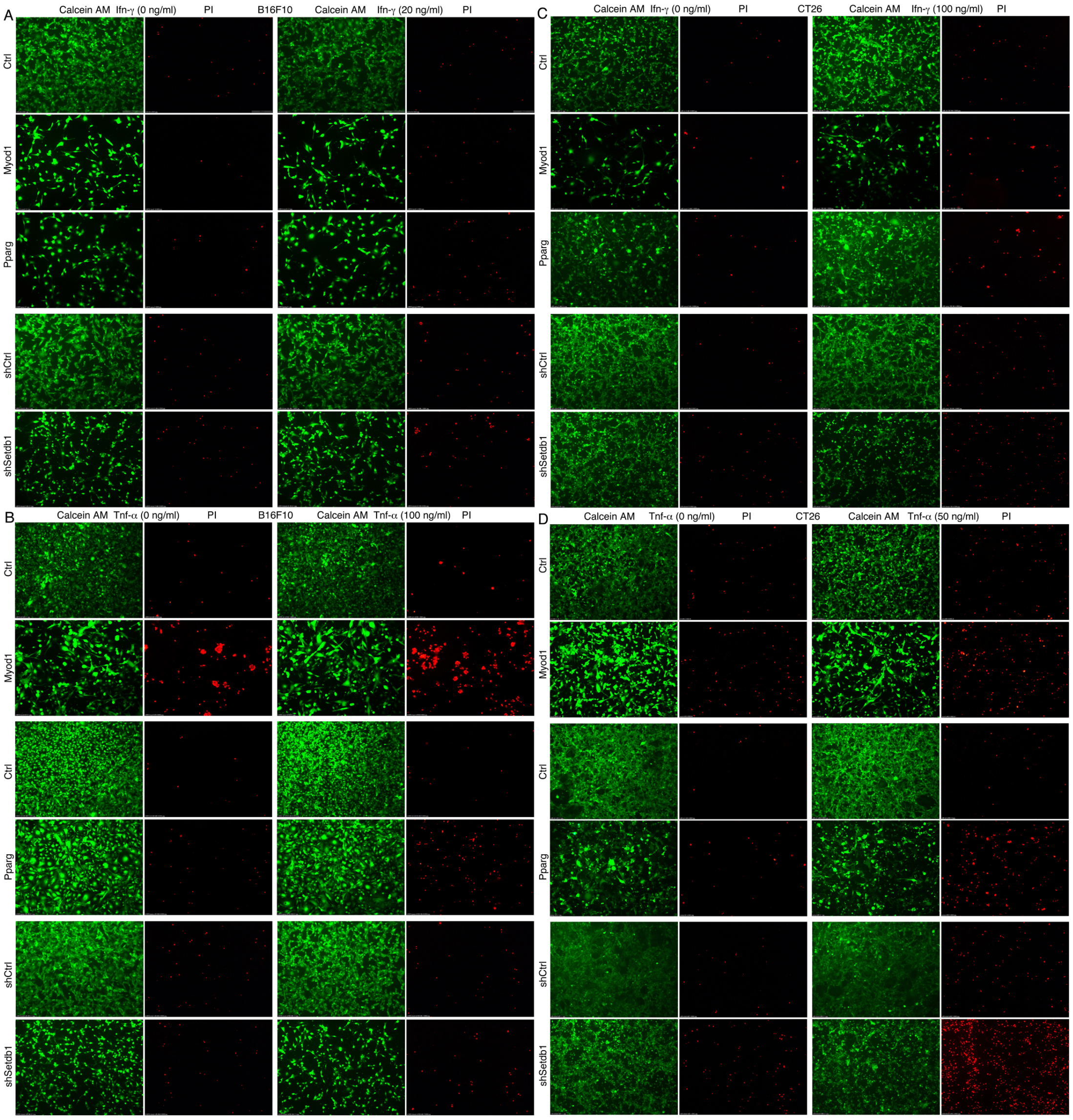


**Fig. S6. Differential response of control and differentiated B16F10 or CT26 cells to induced cell death by Ifn-γ or Tnf-α treatment.** Control or differentiated B16F10 (A, B) or CT26 (C, D) cells induced by forced Myod1 or Pparg expression or knockdown of Setdb1 were subjected to treatment of Ifn-γ (A, C) or Tnf-α (B, D) at indicated doses. Live cells were indicated with Calcein AM staining, and dead cells were indicated with PI (Propidium Iodide) staining, and observed under fluorescence microscope.

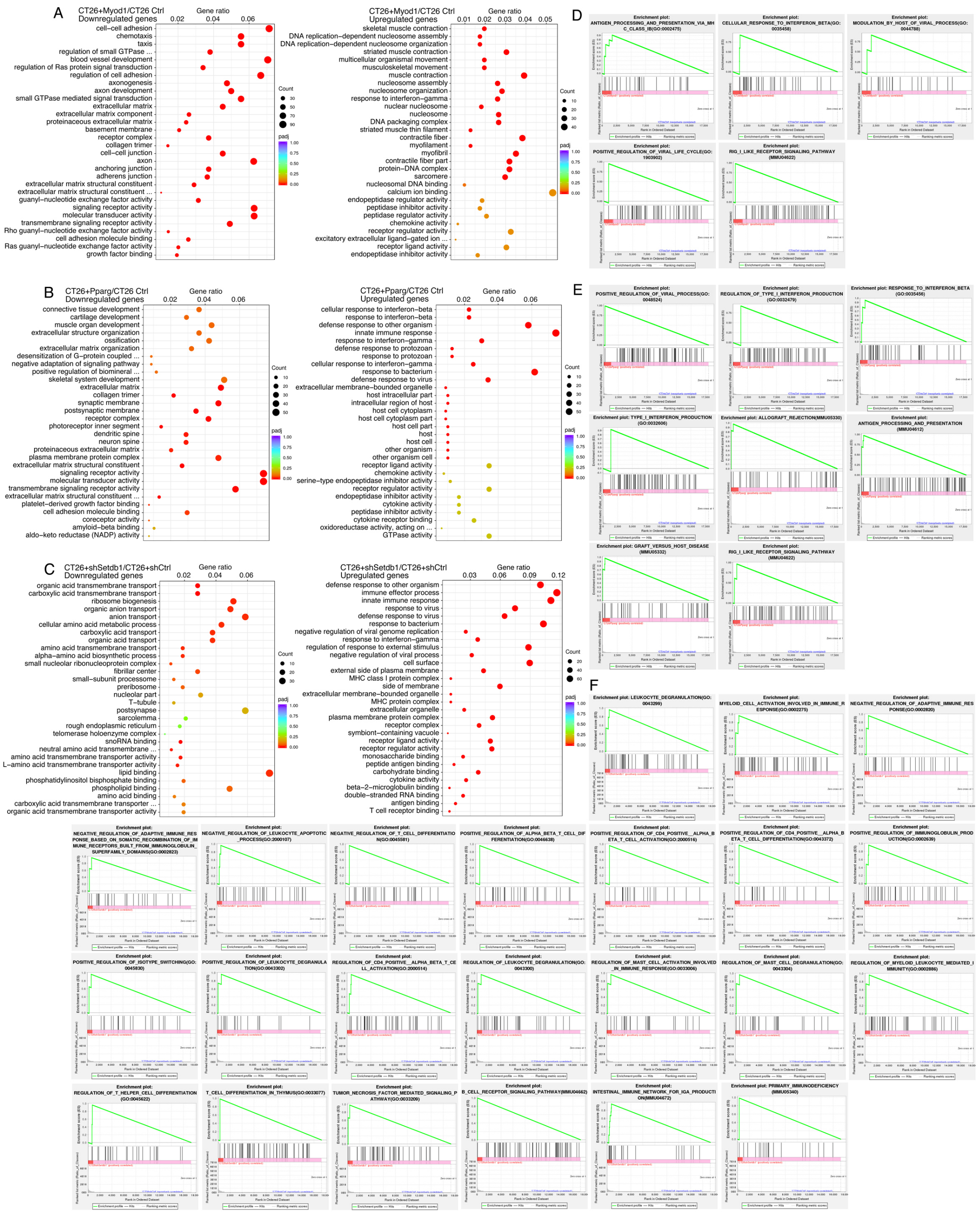


**Fig. S7. Transcriptome profiling of control CT26 cells and cells after induced differentiation.** (A-C) GO analysis on downregulated and upregulated genes in differentiated cells with forced expression of Myod1 (A), Pparg (B), or Setdb1 knockdown (C). (D-F) Immune related gene sets enriched by GSEA in differentiated cells in response to forced expression of Myod1 (D), Pparg (E), or knockdown of Setdb1 (F).

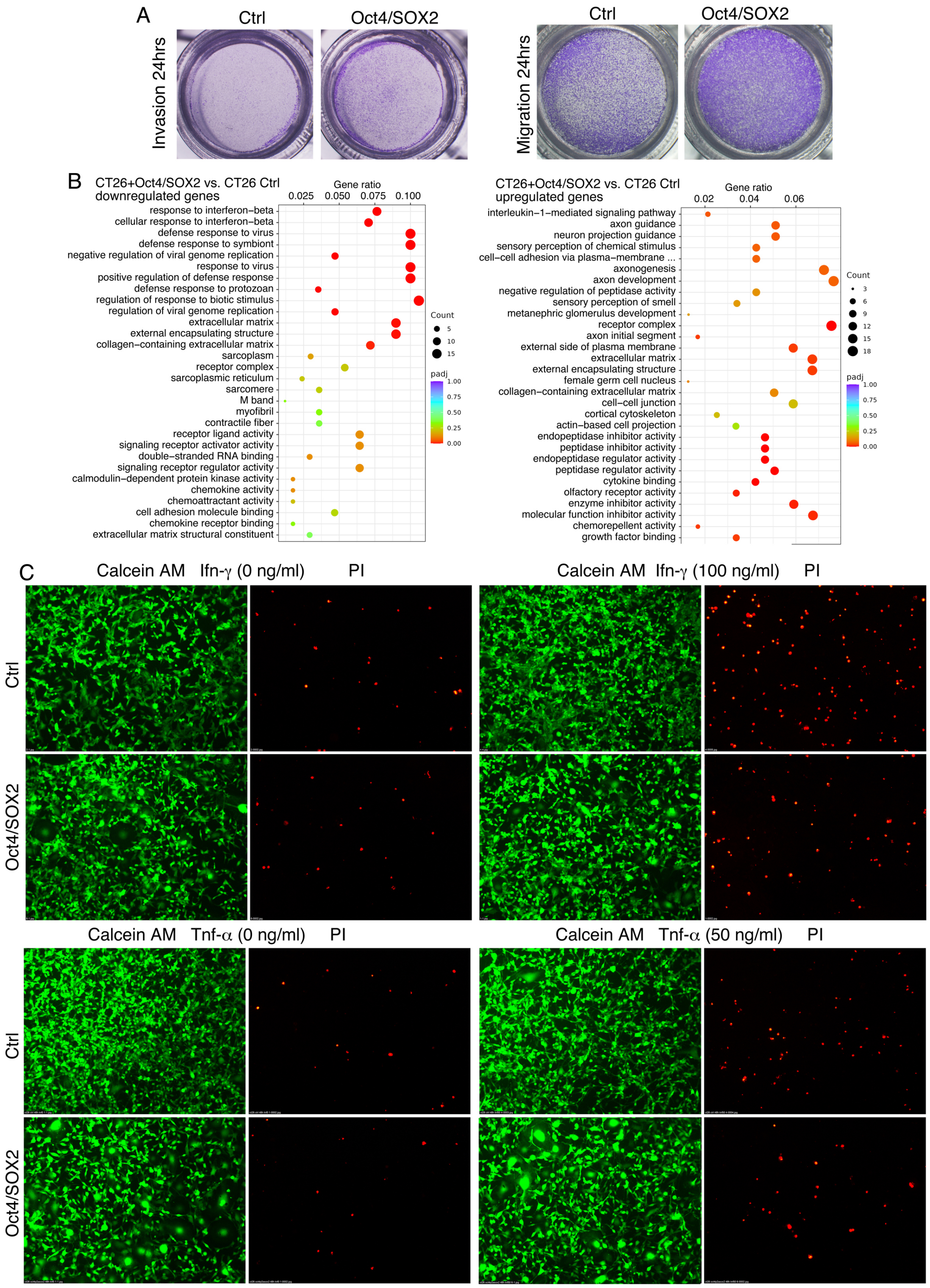


**Fig. S8. Characterization of cellular properties and transcriptomes of control and dedifferentiated CT26 cells by forced expression of Oct4/SOX2.** (A) Invasion and migration assay. (B) GO analysis on differentially expressed genes. (C) Differential response of control and dedifferentiated CT26 cells to induced cell death by Ifn-γ or Tnf-α treatment. Control or dedifferentiated CT26 cells were subjected to treatment of Ifn-γ or Tnf-α at indicated doses. Live cells were indicated with Calcein AM staining, and dead cells were indicated with PI (Propidium Iodide) staining, and observed under fluorescence microscope.

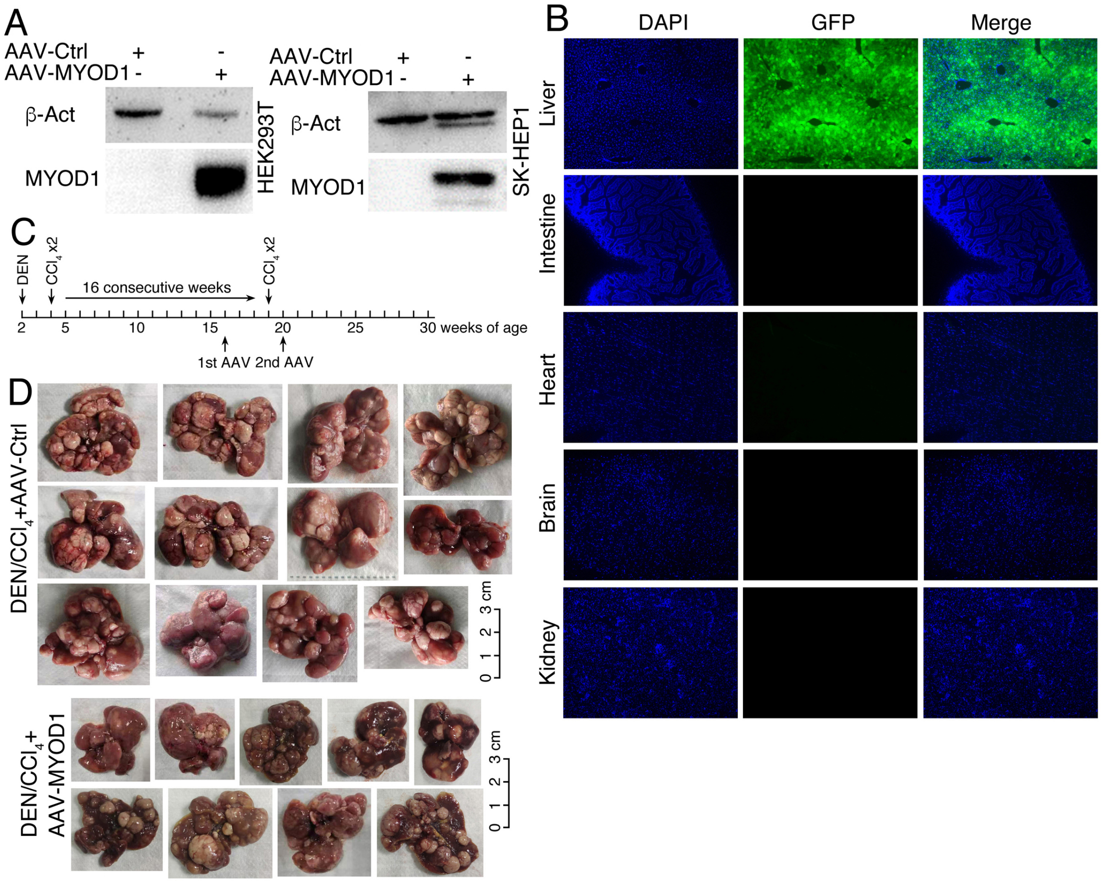


**Fig. S9. Expression effect of AAV-MYOD1 construct and its influence on tumorigenesis in the DEN/$CCl_4$-induced mouse model of hapatocellular carcinoma.** (A) Verification of AAV-MYOD1 construct in HEK293T and SK-HEP1 cells with WB. β-Act was used a loading control. (B) Verification of tissue specific transduction of AAV. Tissues were dissected a month post intravenous injection of AAV. Expression effect was examined by observing green fluorescence in cryosections of different tissues. Nuclei were counterstained with DAPI. (C) Strategy for generation of DEN/$CCl_4$-induced mouse model of hepatocellular carcinoma and time points for AAV injection. (D) Liver tissues from mice with DEN/$CCl_4$ treatment plus injection of control AAV (AAV-Ctrl) and mice with DEN/$CCl_4$ treatment plus injection of AAV carrying Myod1 (AAV-MYOD1).

**Supplemental table legends**

Table S1. Differentially expressed genes involved in IFN-γ response and antigen processing and presentation between control and differentiated HCT116 or SK-HEP1 cells, as revealed by transcriptome profiling.

Table S2. Differentially expressed Ifn-γ response genes, and antigen processing and presentation genes in mouse embryonic stem cells (mESCs), muscle, fat, spleen and thymus cells, and enriched immune related gene sets by GSEA in these cells, as compared with those in mouse primitive neural stem cells (primNSCs) by transcriptome profiling.

Table S3. Differentially expressed genes involved in Ifn-γ response and antigen processing and presentation between control and differentiated B16F10 or CT26 cells, or between control and dedifferentiated CT26 cells, as revealed by transcriptome profiling.
